# Assessing evolutionary and developmental transcriptome dynamics in homologous cell types

**DOI:** 10.1101/2021.02.09.430383

**Authors:** Christian Feregrino, Patrick Tschopp

**Author notes:** Correspondence (CF), (PT).

## Abstract

**Background:** During development, complex organ patterns emerge through the precise temporal and spatial specification of different cell types. On an evolutionary timescale, these patterns can change, resulting in morphological diversification. It is generally believed that homologous anatomical structures are built – largely – by homologous cell types. However, whether a common evolutionary origin of such cell types is always reflected in the conservation of their intrinsic transcriptional specification programs is less clear.

**Results:** Here, using a paradigm of morphological diversification, the tetrapod limb, and singlecell RNA-sequencing data from two distantly related species, chicken and mouse, we assessed the transcriptional dynamics of homologous cell types during embryonic patterning. We developed a user-friendly bioinformatics workflow to detect gene co-expression modules and test for their conservation across developmental stages and species boundaries. Using mouse limb data as reference, we identified 19 gene co-expression modules with varying tissue or cell type-restricted activities. Testing for co-expression conservation revealed modules with high evolutionary turnover, while others seemed maintained – to different degrees, in module make-up, density or connectivity – over developmental and evolutionary timescales.

**Conclusions:** We present an approach to identify evolutionary and developmental dynamics in gene co-expression modules during patterning-relevant stages of homologous cell type specification.

## 1 INTRODUCTION

Recent advances in single-cell technologies now enable researchers to study the molecular dynamics of pattern formation and evolution at the level of the basic biological unit of life, the individual cell. During development, starting from a single fertilized cell, various progenitor cell populations need to proliferate, differentiate, and – for some of their progeny – undergo controlled cell elimination. These processes require tight coordination, across time and space, to result in proper pattern formation of complex organs. From a cell’s perspective, this progression is linked to the integration of various extra-cellular signals, as determined by its relative position inside the forming tissue, and the cell-intrinsic interpretation of these cues, shaped by the lineage-specific molecular state of the cell. Accordingly, evolutionary modifications in a given patterning process can occur through changes in either cell-extrinsic or -intrinsic components – or a combination of both –, to result in morphological diversification.

This is exemplified in the vertebrate limb, a paradigm of morphological evolution, where developmental patterning has experienced important modifications across the tetrapod clade. Molecular genetics studies and experimental embryology have yielded important insights into how, e.g., early limb bud outgrowth is initiated and advanced, or what modifications in these molecular programs can drive diversification of limb form and function.^1, 2^ So far, a majority of these findings have focused on the activity and interplay of various extra-cellular signaling pathways.^3–8^ Yet only since recently, thanks to the development of single-cell genomics technology, can we study the molecular dynamics that occur cell-intrinsically, inside the signal-receiving cell types, at high resolution.

Cell types, like anatomical structures, can be considered homologous across different taxa, with their evolutionary origins tracing back to a common ancestor.^9, 10^ Moreover, it its generally believed that homologous anatomical structures are built – to a large extent – by homologous cell types, and that changes in organ patterning often simply reflect temporal, spatial and quantitative differences in the specification of these cells during development.^11^ The overall molecular state of a homologous cell type, however, may vary substantially between species, even within a similar developmental context. This holds especially true at the transcriptional level, where selection can be weak and result in genetic drift and concerted transcriptome evolution.^12–16^ Accordingly, identification of so-called ‘species signals’, rather than functionally relevant gene expression changes, can dominate differential expression analyses, particularly when applied to similar cell types over long evolutionary distances.^12, 17^ Moreover, any given cell type might occur in a variety of so-called ‘cell states’, related to e.g. cell cycle or metabolic status, thereby further complicating these comparisons.^18^ Hence, to understand developmental pattern evolution comprehensively, we require both a detailed understanding of the cell-extrinsic changes occurring in the signaling environment, as well as appropriate methods to detect species-specific differences in the intrinsic molecular makeup of the recipient cell types.

Here, we present a bioinformatics workflow in *R* to identify gene co-expression modules from scRNA-seq data, and test for their conservation across developmental stages and species boundaries. Using mouse limb E15.5 data as our reference, we identify tissue and cell type-specific co-expression modules and demonstrate the ability to follow their compositional changes, module architecture and expression dynamics along developmental time courses in two distantly related tetrapods. Differences in module conservation – between modules, across species – indicate that patterning processes involving certain cell populations are more likely to occur through changes in extracellular environment, while others undergo high evolutionary turnover in their cell-intrinsic molecular make-up. Moreover, we demonstrate the power of gene co-expression module detection to identify distinct cell states, shared across developmental and evolutionary timescales.

## 2 RESULTS

### 2.1 Primary data acquisition, 3’UTR annotation and data processing

We used publicly available scRNA-seq data from mouse and chicken spanning six embryonic stages, from early limb bud initiation and outgrowth, to late stages of pattern refinement and tissue maturation. For mouse, we had access to stages E9.5, E10.5, E11.5, E13.5, E15.5, and E18.5, and used a total of 17857 cells^19, 20^ (Fig. 1A). For chicken, we used our own previously published data (HH25, HH29, and HH31^21^), and complemented the time series with newly generated data points spanning stages HH21, HH24, and HH27 (Fig. S1A, B), to cover days 3 to 7 of development with a total of 32461 cells (Fig. 1B). For both chicken and mouse, we have fore- and hindlimb data, which we analyzed interchangeably (Fig. 1A, B).

**Figure 1.**
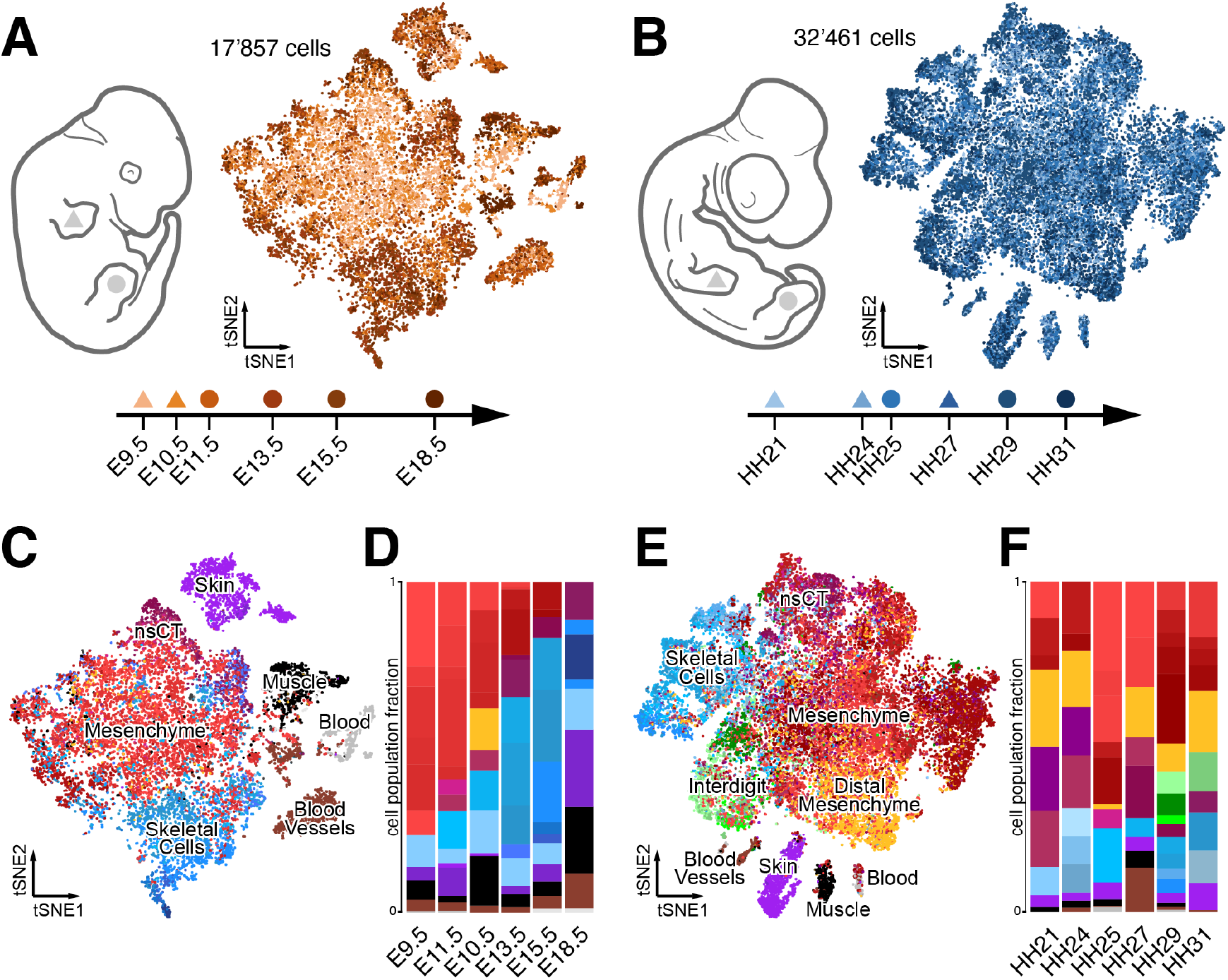
Comparative single-cell transcriptomic atlases of the developing mouse and chicken limb. (A, B) Sampling schemes and tSNE projections of 17’857 mouse (A) and 32’461 chicken (B) forelimb (triangle) and hindlimb (circle) single cell transcriptomes. (C-F) Mouse (C) and chicken (E) cluster identities are highlighted by color codes in the tSNE projections, on a cell-by-cell basis, with cells belonging to similar tissue types sharing color codes across all samples. Relative sample composition is visualized by barplots (D, F). A total of 86 and 74 clusters were identified in mouse and chicken, respectively, distributed over 6 sampling time points.

A preliminary inspection of the two data sets revealed that – on average – mouse samples displayed a higher percentage of skeletal cell types. We reasoned that either during the preparation of single cell suspensions the mouse tissue had been dissociated more thoroughly, thereby releasing a higher percentage of extracellular matrix-encapsulated skeletal cells, or that our expression analyses of chick cells failed to accurately capture the expression status of genes important for skeletogenesis. Indeed, when visually examining the genomic location of our mapped chicken reads, many seemed to fall outside the annotated 3’ UTR regions and hence were not included in our UMI count tables (Fig. S1A, B). Such annotation issues have been reported before, also for other species,^22^ and they are particularly problematic when using sequencing technologies with high 3’UTR-biases like the *10x* Genomics Chromium 3’ Kit. Therefore, we decided to improve the 3’UTR annotation of the chicken genome, using publicly available bulk RNA-seq data (see Experimental Procedures).^23^ This resulted in a slight overall increase of the average 3’UTR length, yet with many of the extensions not exceeding 100bp (Fig. S1C, D). However, even such modest extensions in 3’UTR lengths resulted in a substantial increase of UMI numbers detected for many genes, including some well-known skeletal regulators like *SOX9* and *GDF5* (Fig. S1E, F). Accordingly, we re-mapped all our chicken data using our newly improved 3’UTR annotation. While this did not completely alleviate the bias in murine skeletal cells – i.e. differences in dissociation protocols likely also contribute to this effect –, we now managed to identify small skeletal sub-populations, like e.g. synovial joints, more reliably in the chicken (data not shown). Moreover, we believe that this improved 3’UTR annotation will also prove helpful for future single cell genomics studies in the chicken model system.

With these improved UMI count tables for the chicken, we continued our comparisons to the mouse limb samples. As our datasets were produced in different laboratories, we implemented a standardized filtering step of all single cells, based on quality measurements like library size, proportion of mitochondrial reads and number of genes detected. Moreover, due to the overall size of the E9.5 and E10.5 datasets, we randomly subsampled 25% of the single cell transcriptomes, to have datasets of comparable sizes. We then normalized the expression data, performed cell cycle correction and adjusted multi-batch samples. For each species, we then integrated all cells into a single tSNE dimensionality reduction embedding (Fig 1A, B).

In order to identify the different cell types in our data, we first analyzed all stages individually. Using the same parameters of unsupervised graph-based clustering throughout, we found 13 clusters in the mouse E9.5 sample, 16 at E10.5, 12 at E11.5, and 15 at E13.5, E15.5 and E18.5 each. For our chicken samples, we found 9 clusters in the HH21 dataset, 11 at HH24, 15 at HH25, 9 at HH27, 19 at HH29, and 11 at HH31. By comparing the results of our differential gene expression analyses to known marker genes, we were able to identify most of these clusters as distinct cell or tissue types. In all samples we found one major cell population, consisting of lateral plate mesoderm-derived limb mesenchymal cells at various stages of differentiation, as well as several smaller clusters of cells with different developmental origins (Fig 1C, E). Of those, skin cells (purple) were present in all samples, while muscle cells (black), blood (light gray) and blood vessels (brown) were detected only in a subset of the samples. A small cluster of likely melanocytes (dark gray) was found only at mouse stages E15.5 and E18.5.

Within the lateral plate mesoderm-derived limb mesenchymal cells we identified undifferentiated limb mesenchyme (light red), proliferating or cycling mesenchyme (dark red), non-skeletal connective tissue (nsCT) (maroon), and skeletogenic cells like e.g. chondrocytes (blue). Mesenchymal cells with a likely distal origin (yellow) were detected only at stage E11.5 in the mouse, but in all chicken samples, while the interdigit cells were only found in the later chicken stages HH29 and HH31 (green). Differences in dissection strategies and dissociation protocols likely account for these disparities in cell types detected in a given sample, as well as for changes in their relative abundance. For example, while in mouse samples coming from whole limbs a steady increase in the proportion of skeletal cells is observed, a more targeted sampling of certain limb sub-domains in the older chicken samples likely obscured this effect (see ref.^21^ for details) (Fig 1D, F). Overall, in 17857 mouse cells and 32461 chicken cells we identified a total of 86 and 74 clusters, many of which correspond to distinct cell types at various stages of maturation across the 6 developmental stages. More importantly, the large majority of these cell types can be considered homologous between the two species, and hence their single-cell transcriptomes can now be used in a comparative context, to assess cell type-specific transcriptional dynamics across developmental and evolutionary timescales.

### 2.2 A bioinformatics workflow to detect cell type-specific gene coexpression modules from scRNA-seq data

To detect cell type-specific gene expression signatures, and circumvent some of the issues inherent to crossspecies differential expression analyses, we adapted weighted gene correlation network analysis (WGCNA)^24^ and tested for the occurrence of transcriptome-wide gene co-expression patterns in single cells. WGCNA, originally developed for the detection of gene co-expression modules in bulk RNA-seq data, has seen a recent surge in popularity, given the high number of replicate samples, i.e. single cells, available when working with scRNA-seq data. We reasoned that a standardized, user-friendly bioinformatics workflow would proof beneficial to first-time users of WGCNA, as well as make the results more comparable between different types of studies and data sets. Accordingly, we developed an *R* package (“scWGCNA”) for gene co-expression module detection and comparisons. In a first part, our analysis starts with a Seurat object^25^ – one of the most commonly used output formats of scRNA-seq data analyses these days – and then performs, 1) pseudocell construction, to increase overall robustness; 2) identification of highly variable genes [if not already provided by the user]; 3) WGCNA module detection; 4) Gene Ontology (GO)-term enrichment analyses and putative cell type identification; and, lastly, produces 5) a standardized output file in HTML format.

For pseudocell construction, 20% of the cells from each cluster are chosen at random (see Fig 2A), to which their 10 nearest neighboring cells in the PCA space are then aggregated.^26^ The average expression of every gene is calculated for each of these cell aggregates and normalized, to result in a gene-by-pseudocell expression data matrix. We consider several metadata bins contained in the original Seurat object, including – if already calculated – a set of highly variable genes as determined from the single-cell data. This gene set is critical for subsequent analyses, as it directly affects the modules that potentially can be detected.^24, 27^ Accordingly, ‘highly variable gene detection’ is optional in our pipeline (see above), allowing users to opt for their method of choice. Thereafter, using the variable genes expression matrix as input, a range of powers are tested to find a suitable soft-thresholding power that transforms the correlation network to resemble a scale-free topology – i.e. the underlying structure and characteristics are independent of changes in network size, which is assumed to be the case for biological networks.^24, 28^ This step is inherent to WGCNA and aims to reduce the noise of correlations in the adjacency matrices used. Moreover, it also serves as an important control point: if a scale-free topology index is not reached, the genes – or cells – used should be reconsidered by the user. Next, the main WGCNA analysis follows. In short: based on the expression matrix, expression correlation, adjacency, and topological overlap matrices are calculated first. Then, based on topological overlap, genes are assigned to discrete modules of co-expression. The membership of the genes to their modules is tested, based on the correlation of their expression to the overall expression of the module: genes without significant membership are discarded, and the process is repeated until all genes pass the membership test. Lastly, the mean expression of the different modules is calculated in single-cell space, and plotted onto a dimensionality reduction of choice. Graph representations of all modules are generated, and corresponding GO-term enrichment analyses are performed. All results are then summarized in a single report in HTML format (see Supplement 1). We decided to test our workflow with only one limb data set, in order to be able to compare composition and expression status of the identified modules in different embryonic stages, as well as across species boundaries. We used the mouse E15.5 sample as our reference data, as it showed a high variety in skeletal and connective tissue cell populations (Fig. 2A).

**Figure 2.**
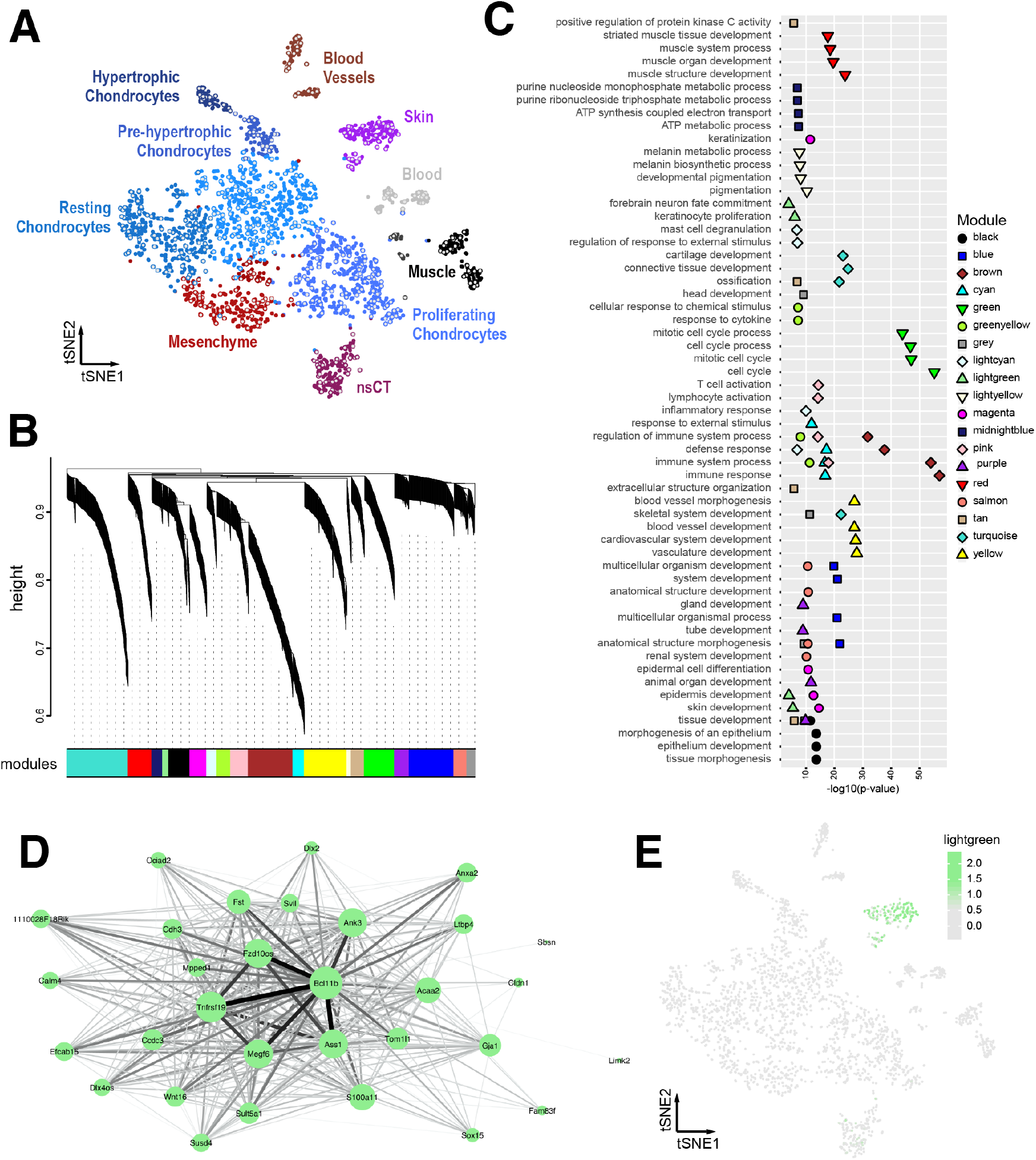
A bioinformatics workflow for iterative gene co-expression module detection and identification. (A) tSNE projection of the mouse E15.5 sample used as reference for iterative WGCNA gene co-expression module detection. Identified clusters are color-coded and labeled, with seed cells used for pseudocell construction highlighted in grey. (B-E) Sample outputs of our ‘scWGCNA’ package. (B) Final WGCNA gene hierarchical clustering dendrogram, showing 19 modules of coexpression. (C) Cumulative results of GO-term enrichment analyses, showing the top 4 (by p-value) enriched GO-terms for each module. (D) *Cytoscape* visualization of co-expression module ‘lightgreen’. Node size for each gene is proportional to its module membership, edge thickness and intensity represent topological overlap. (E) E15.5 tSNE showing the averaged cellular expression of module ‘lightgreen’.

From the 2594 E15.5 single-cell transcriptomes a total of 513 pseudocells were constructed. Using the 2967 top variable genes as input, we obtained 1248 genes showing significant co-expression dynamics, distributed over 19 modules of co-expression. For sake of simplicity, the different modules identified are hereafter referred to by their colors, according to the WGCNA results (Fig. 2B, C). The modules varied substantially in size and degrees of similarity among them (Fig. 2B). Genes within the detected modules showed signs of enrichment for functional GO-terms that we expect to be important for limb development, distributed over the different tissue types found in the forming appendage (Fig. 2C). Likewise, the averaged expression of a module, calculated over all single cells as the averaged expression of all the genes contained within it, often showed patterns of tissue or cell type specificity. Certain modules exhibited highly restricted expression, confined to a single cell cluster, while others spanned across multiple populations (Supplement 1). For example, module ‘lightgreen’ was one of the smallest modules detected, with 31 genes centered around *Bcl11b*, and an averaged module activity confined to our previously identified skin cluster (Fig. 2D, E). Other modules with tissue specificity and indicative GO terms were, e.g., ‘red’ (muscle), ‘yellow’ (blood vessels) or ‘turquoise’ (cartilage); while modules ‘green’ and ‘midnightblue’ displayed broader patterns of activity and GO term enrichments for “cell cycle” and “cellular respiration”, respectively (Supplement 1).

Hence, our analysis revealed the existence of several gene co-expression modules in one of the limb scRNA-seq data sets, mouse E15.5, with varying patterns of tissue or cell type specificity.

### 2.3 Testing for gene co-expression module conservation across developmental and evolutionary timescales

We then conducted a comparison of these modules of gene co-expression across all samples, testing for the conservation of different properties in each module: gene composition, ‘density’ and ‘connectivity’. First, to assess module composition and overall activity between the two species, we checked if genes within the modules were present as orthologues, and whether they are expressed in any of the chicken samples. In terms of gene content, we only considered 1-to-1 orthologues, based on *Ensembl* criteria with a confidence cut-off of 1.^29, 30^ Presence/absence of genes in the chicken genome varied greatly between the different modules, with the highest percentage of 1-to-1 orthologues missing in modules related to immune function (Fig. 3A, ‘brown’, ‘greenyellow’, ‘pink’, ‘lightcyan’ and ‘cyan’). Conversely, modules enriched for GO-terms related to transcriptional regulation and morphogenesis all had more than 75% of their genes represented in the chick genome (Fig. 3A, ‘purple’; and ‘blue, ‘salmon’). In terms of expression, the highest fraction of non-expressed genes was found in modules ‘magenta’ and ‘lightyellow’, with ~15% and ~25% of their 1-1 orthologous not being detected in our chicken samples. Overall, our analysis showed that for 8 of the 19 modules less than 60% of their genes are present in our chicken samples as 1-to-1 expressed orthologues. Seven of these 8 modules were enriched for GO-terms related to immune function or skin development (Fig. 3A and Supplement 2). For the remainder of our analyses, we only considered 1-to-1 orthologues expressed in samples of both species.

**Figure 3.**
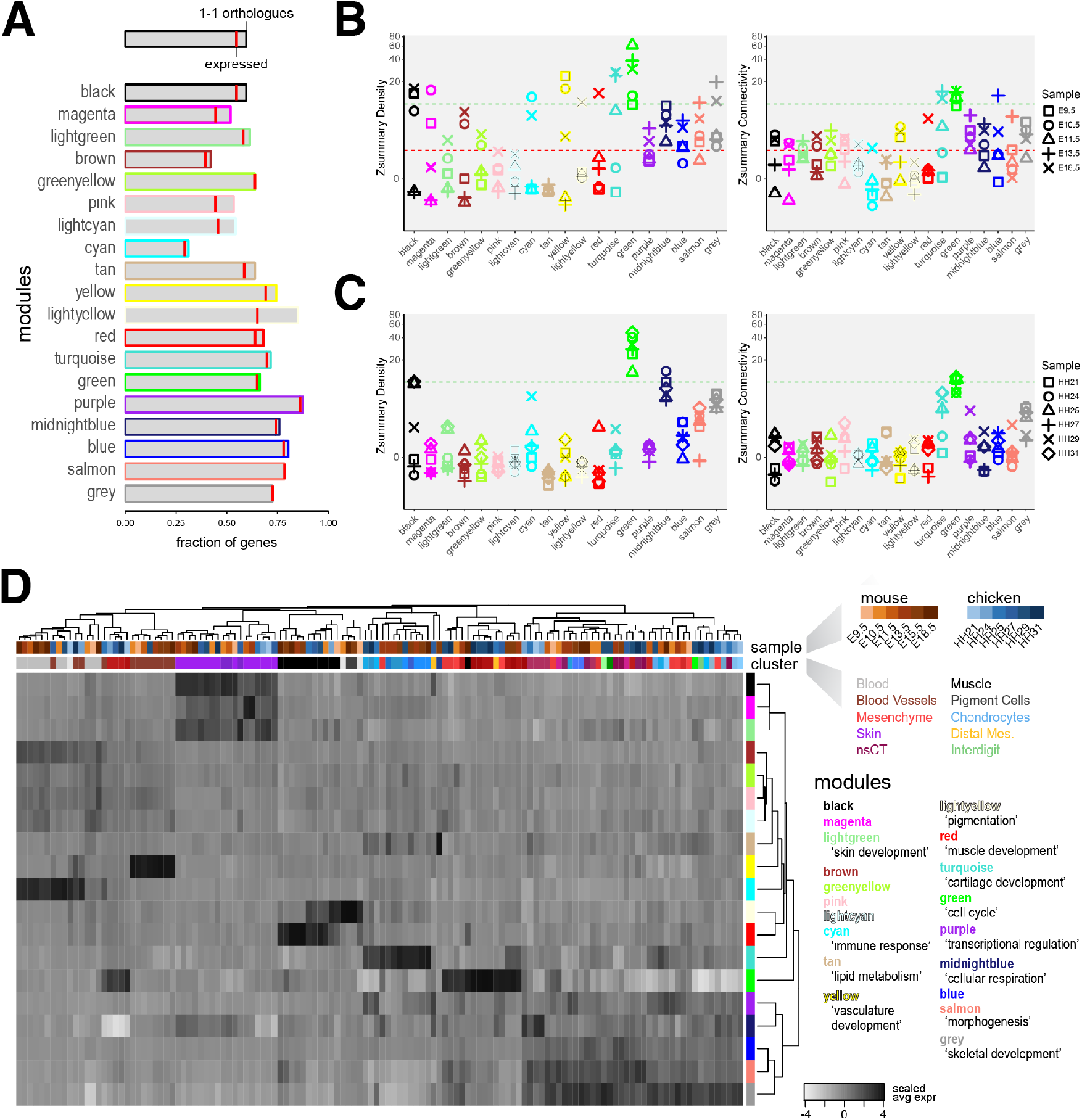
Evolutionary and developmental dynamics in gene co-expression modules. (A) Conservation of module gene composition and expression status. Barplots showing, module by module, the fraction of module genes present as 1-1 orthologues in the chicken genome, and whether or not they are expressed in our samples (red bar). (B, C) Zsummary statistics for ‘density’ and ‘connectivity’, module by module, for each test sample in mice (B) and chicken (C). The different developmental stages for each sample are coded for by symbols. Green and red dotted lines indicate Zsummary values of 10 and 2, respectively, to indicate ‘strong evidence’ of module ‘density’ or ‘connectivty’ conservation (above 10), potential conservation (between) or ‘no evidence’ of conservation (below 2). (D) Hierarchically clustered heatmap of scaled averaged expression for each gene co-expression module, across mouse and chicken samples on a cluster-by-cluster pseudobulk basis. Top row of cluster identifiers is color-coded for species and developmental stage, bottom row for tissue type to which the respective cluster was attributed to, based on marker gene expression. Representative GO-terms are provided for each module.

We next wanted to assess to what extent the module co-expression relationships between genes are conserved across developmental and evolutionary timescales. A co-expression network can be visualized as a group of genes (nodes), connected with different strengths as defined by their co-expression relationships (edges) (see also Fig. 2D). We tested for conservation of ‘density’ (i.e., the average strength of all connections between all genes) and ‘connectivity’ (i.e., the patterns of strength of connections) in all of our modules.^31^ For this, we developed a second part of our workflow, again implemented in our *R* package. As input, a list of 1-to-1 orthologous genes, expression matrices of the test datasets, and the reference WGCNA analysis (see above) is required. In a first step, modules are filtered to only contain expressed 1-to-1 orthologues. In a second step, a “preservation test” is performed, which integrates a suite of statistical tests of network properties into so-called “preservation indices”.^31^ Hence, we obtain metrics for the conservation of module gene composition, as well as indices for the preservation of ‘density’ and ‘connectivity’ for each module, across the different test samples (Fig. 3A-C, Supplement 2).

These “preservation tests” showed that the co-expression dynamics within our modules have different levels of conservation across our samples. To quantify these differences, we used the integrated Zsummary statistic of conservation. This statistic can be interpreted with two thresholds, with a Zsummary greater than 10 implying strong evidence of module preservation, and lower than 2 suggesting no evidence of preservation.^31^ In general, we observed that – as expected – the co-expression relationships of the modules detected in the E15.5 sample are more conserved in the other mouse samples, than in chicken. We also found that, overall, density is more conserved than connectivity, with only a few exceptions (Fig. 3B, C). By using a median rank index, we observed that the most conserved modules are ‘green’ (GO-term enrichment (GOE): ‘cell cycle’) and ‘grey’ (GOE: ‘skeletal development’). In the mouse samples, we noticed that only module ‘purple’ (GOE: ‘transcriptional regulation’) shows an overall higher conservation of connectivity than density, implying that the co-expression relationships between specific genes are better conserved than the overall correlation in the module. Moreover, in chicken samples, modules ‘purple’ and ‘turquoise’ (GOE: ‘cartilage development’) also showed overall higher conservation of connectivity than density. On average, however, modules related to ‘cartilage’ and ‘skeletal development’ showed higher conservation in density and/or connectivity in chicken samples, as compared to module ‘purple’, even though the latter contained a higher fraction of expressed 1- to-1 orthologues (Fig. 3A-C, Supplement 2).

Despite this seemingly low level of overall conservation, our gene co-expression modules still seemed to carry a substantial amount of information concerning cell type and cell state, both across developmental stages as well as for comparing samples between the two species. To try and infer cell type and cell state equivalencies, we first calculated the expression of each module for every cell. We then averaged cellular module expression across all previously identified cell clusters, for each of our samples, to define so-called cell cluster-specific “pseudobulk” representations of module activities. Importantly, our modules calculated from the mouse E15.5 sample might not accurately reflect the transcriptional activity of all cells in our study. To account for this, we defined a threshold as the median expression of all modules in a given pseudobulk, plus two times the median absolute deviation. Only pseudobulks expressing any module at a higher level than this threshold were considered for further comparisons. Out of a total of 160 pseudobulks, 128 showed high enough expression of at least one of the co-expression modules to pass our threshold. We scaled the expression data module-wise and calculated Pearson’s correlation coefficients, Euclidian distances and hierarchical clustering of all pseudobulks and modules. For the most part, these pseudobulks didn’t group by species in the hierarchical clustering, but rather by cell type in general (Fig. 3D). Pseudobulks not derived from lateral plate mesoderm cells showed a particularly clear cell type-based clustering, regardless of the species of origin. We found chicken and mouse blood cells, vessels, muscle and skin pseudobulks grouped together, due to their elevated expression of modules enriched for GO-terms reflecting the respective cellular functions. On the other hand, lateral plate mesoderm-derivatives were divided into three major clusters, based largely on the expression of modules ‘turquoise’ (GOE: ‘cartilage development’), ‘green’ (GOE: ‘cell cycle’), and ‘purple’ to ‘grey’ (GOE: ‘transcriptional regulation’, ‘cellular respiration’, ‘morphogenesis’ and ‘skeletal development’) (Fig. 3D).

Collectively, testing for conservation of gene co-expression at multiple embryonic timepoints, and between distantly related species, revealed considerable disparities between the different modules. Often, these differences were in line with the likely cellular functions attributed to the respective modules. Regardless of the degree and type of conservation, however, most of the identified modules still seemed to contain important information concerning the cell type and state from which a given single-cell transcriptome originated from, both across different developmental stages and taxa.

### 2.4 Cross-species developmental dynamics and ontogenetic trajectories of gene co-expression modules

Finally, we analyzed the expression of our identified modules across embryonic time, in both species, taking advantage of the different developmental stages that were used for tissue sampling. We focused only on cells derived from the early limb bud mesenchyme, as they have a common developmental origin in the lateral plate mesoderm, play a central role in establishing the eventual limb morphology, and displayed a higher degree of heterogeneity in module activities amongst themselves (see Fig. 3D). We selected modules showing high scaled averaged expression in these cells, and all lateral plate mesoderm-derived pseudobulks as input. In order to appreciate developmental changes in module gene expression, we re-grouped the corresponding cell cluster pseudobulks, species by species, by computing pair-wise Pearson’s correlation coefficients, Euclidian distances and hierarchical clustering. Based on tree height, this identified 4 major clusters for the mouse and 5 for the chicken. For both species, we additionally identified 2 clusters, each consisting of only two pseudobulks, which we chose not to analyze further (Fig 4A, B). These module-defined clusters roughly equated to ‘mesenchyme’, ‘proliferative mesenchyme’, ‘nsCT’ and ‘chondrocytes’, when comparing them to the original assignments of their respective cell cluster pseudobulks (Fig 4A, B).

**Figure 4.**
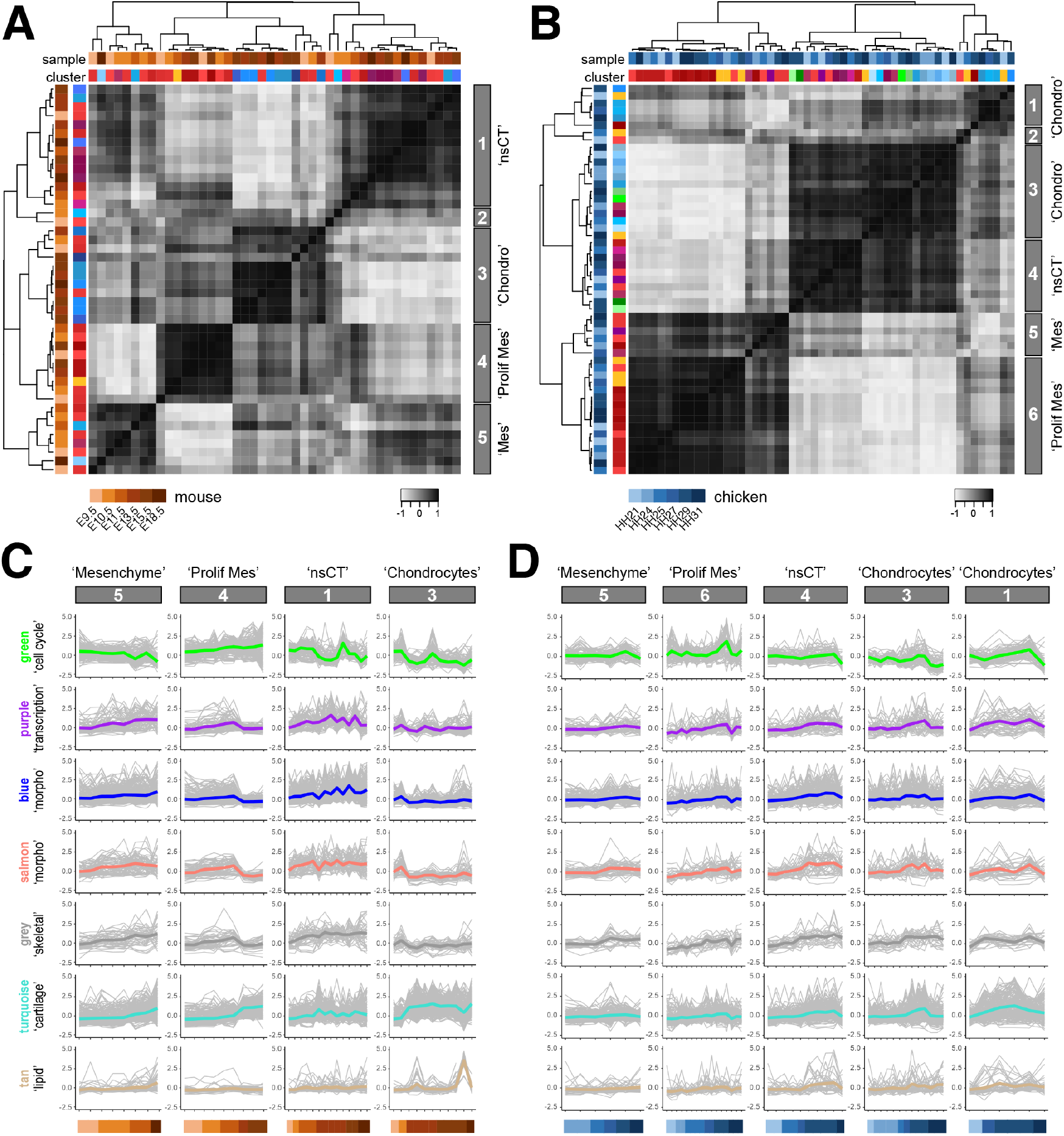
Comparative developmental transcriptome trajectories in homologous cell populations. (A, B) Heat maps of Pearson’s correlation coefficients and hierarchical clustering based on pairwise Euclidian distances, calculated on averaged expression of selected modules, for mouse (A) and chicken (B) lateral plate mesoderm-derived cell cluster pseudobulks. Top row of cluster identifiers is color-coded for species and developmental stage, bottom row for tissue type to which the respective cluster was attributed to, based on marker gene expression. The five (A, mouse) and six (B, chicken) major clusters emerging, based on tree height, are highlighted on the right with grey boxes. (C, D) Developmental trajectories of scaled averaged module expression in ontogenetically ordered pseudobulks, separated by ‘tissue-like’ clusters identified in A, B. Embryonic stage of each pseudobulk is color-coded at the bottom. Scaled averaged modules expression trajectories shown in the respective module color, with induvial module gene traces underlaid in grey.

For mouse and chicken, within each module-defined cluster, we ordered the pseudobulks according to their embryonic stage of collection and plotted the scaled averaged module expression along these ontogenetic trajectories. Additionally, we included the individual gene expression traces represented in the respective module activities, (Fig 4C, D). We observed that the genes in module ‘green’ (GOE: ‘cell cycle’) increased their expression along development only in the ‘proliferative mesenchyme’, while they decreased in ‘nsCT’ and ‘chondrocytes’. Module ‘purple’ (GOE: ‘transcriptional regulation’) showed no reproducible trend in activity, between the different clusters and the two species. Genes in modules ‘blue’, ‘salmon’ and ‘grey’ (GOE: ‘morphogenesis’ and ‘skeletal development’) all showed very similar dynamics, with increased expression along the development of the ‘nsCT’ cluster in both species. Module ‘turquoise’ (GOE: ‘cartilage development’) increased its activity in the ‘chondrocyte’ clusters of mouse and chicken, right around the expected onset of chondrogenesis in the two appendages, albeit with high variability in the individual gene expression traces. Finally, module ‘tan’ (GOE: ‘lipid metabolism’) only showed signs of high expression in the hypertrophic cell population in mouse of the E15.5 sample. (Fig 4C).

## 3 DISCUSSION

To understand the molecular basis of morphological evolution, as driven by changes in embryonic and post-embryonic development, both cell-extrinsic and – intrinsic alterations need to be considered.^32^ Here, we present an integrative approach to perform comparative gene co-expression analyses at the single-cell level. We demonstrate its functionality by testing single-cell transcriptomic data from the developing mouse limb for the occurrence of cell type-specific gene co-expression modules and assess their conservation and developmental dynamics in the corresponding cell populations of the chicken.

### 3.1 Assessing gene co-expression modules in scRNA-seq data from distantly related species

Deciphering species-specific molecular states of homologous cell types is essential, to correctly interpret their response to alterations in extracellular signaling environments. With the advent of single-cell genomics, we now have the technological means to perform such analyses at the appropriate cellular resolution, across different species.^33^ However, comparing gene expression between distantly related taxa has its challenges, especially when working with sparse data like scRNA-seq.^12,22,34,35^ To circumvent some of these inherent issues, we decided to test for the dynamics and conservation of gene co-expression modules in pseudocells.^36–39^

In a first step, we follow the logic of an iterative approach, to perform and optimize WGCNA gene co-expression modules calculations within a reference scRNA-seq data set of choice. We use WGCNA statistics to measure significance of gene membership to their assigned modules, and re-group them accordingly for successive rounds of clustering and testing.^21,40,41^ It is important to note here that WGCNA does not reveal *de facto* regulatory networks or functional relationships between genes, but rather simply reflects modules of gene co-expression.^24^ For example, while the coexpression of transcription factors and their putative target genes might indeed reflect regulatory interactions, relying on gene expression data alone to infer this process is prone to result in a high proportion of false positives.^42^ Alternative approaches, making use of properly annotated *cis*-regulatory sequence information, may seem more appropriate for such purposes.^43, 44^ However, the application of such algorithms is mostly restricted to a very limited set of model species, as they rely on the availability of extensive and high-quality transcription factor binding motif data sets.

Accordingly, in the second step of our workflow, we opted to perform comparative analyses using modules of gene co-expression, to make it applicable to the largest number of species possible. Within these modules, we specifically tested for the preservation of the overall strength of connections, i.e. ‘density’, as well as for the patterns of those connections between genes, i.e. ‘connectivity’.^31^ The validity of such comparisons obviously depends on the presence of corresponding cell populations between the samples, as well as the number of orthologous genes found in each species to be compared. Naturally, detection of true 1-to-1 orthologues is bound to decrease with increasing evolutionary distance.^45, 46^ On a module-by-module basis, however, differences in this overall trend may be informative in itself, to interpret the underlying evolutionary dynamics (see Fig. 3A, and discussion, below). As for homologous cell types, the restricted presence – e.g., hypertrophic chondrocytes in the mouse E15.5 sample – or absence – e.g., distal mesenchyme – of certain cell populations in the reference data set also has implications for our comparative analyses (Fig. 2A, B). For example, imagine a gene with strong topological overlap to a cell type-specific module in the reference sample. If that gene in the test sample is co-expressed with different genes in an additional cell population – i. e. absent from the reference sample –, then this might skew connectivity of the tested module. We therefore advise for an informed and balanced selection of the cell populations considered, in order to obtain the most meaningful results.

Importantly, the entirety of the workflow presented above is wrapped in an *R* package with functions that can be run independently, are customizable, and produce standardized output files to serve as input for further in-depth analyses. All necessary code and documentation are publicly available.

### 3.2 Conservation of gene coexpression modules in the developing tetrapod limb

Working with mouse limb E15.5 data as our reference, we identified a total of 19 gene co-expression modules and tested for their conservation in mouse and chicken samples, at multiple developmental timepoints. Already at the compositional level, important qualitative and quantitative differences emerged between the modules. For example, among modules enriched for immune functions, some showed as few as 30% of their genes to be present as 1-to-1 orthologues in the chicken genome (Fig. 3A). Such high genomic turnover is considered a hallmark of the immune system, compared to other functional groups of genes, as it constantly adapts in an evolutionary arms race to an everchanging pathogen and parasite regime.^47, 48^ Likewise, modules enriched for skin-related functions showed low levels of compositional conservation. The function of the skin, and its associated ectodermal appendages (i.e. hair follicles, glands or feathers), has diverged considerably between mammals and sauropsids.^49–51^ Moreover, selection for a variety of integumentary traits in domesticated chickens might have accentuated this trend further.^52^

In terms of ‘density’ and ‘connectivity’, module ‘green’ showed the overall highest degree of conservation, both for mouse and chicken samples (Fig. 3 B, C, Supplement 2). This is somewhat expected, as a coexpression module reflecting the cell cycle process likely should be conserved even between distantly related taxa. For modules predominantly active in lateral plate mesoderm derivatives, certain tendencies emerged when comparing them across developmental time. Overall, ‘density’ and ‘connectivity’ of these modules seemed better conserved in samples at later stages of development (Fig. 3 B, C). Likewise, we observed that early pseudobulks of less differentiated cell populations were underrepresented in our analysis of module expression levels (Fig. 3 D). 9 out of the 14 excluded pseudobulks in mouse, and 6 out of 18 in chicken, stem from mesenchyme populations of our earliest two time points. Moreover, at the finer scale of our hierarchical clustering, pseudobulks from earlier stages tend to cluster by species (Fig. 3 D). The fact that we calculated our reference modules at a rather late stage of development might potentially explain this tendency, i.e. the expression of certain modules might simply not be adequately represented in these early cells. However, by recreating the same analysis using E11.5 modules as reference, we observed a similar trend (data not shown). Therefore, we suggest that advanced differentiation of cell types effectively makes them – at least module-wise – transcriptionally more similar to their counterparts in other species, than to their less differentiated relatives in the same organism.^10, 53^

Of all the modules identified for a distinct cell or tissue type, ‘turquoise’ (GOE: ‘cartilage development’) was the overall largest and showed the highest degree of conservation, (Fig. 2A, Fig. 3A-C). This was particularly evident at later stages of development, and for Zsummary ‘connectivity’, implicating that differentiating chondrocytes indeed follow similar molecular programs in the two species. Specifically, this evolutionary conserved ‘connectivity’ indicates that genes of module ‘turquoise’ share conserved coexpression dynamics, or that they are controlled by the same up-stream factor(s) across taxa. The cells producing the cartilage template of the limb skeleton thus seem equipped with a similar molecular make-up, hence making patterning changes between species likely to occur predominately through alterations in extracellular signaling. However, not all signal-receiving cell populations of patterning relevance show equal conservation in their gene co-expression dynamics. Modules related to skin development show, as outlined above, high compositional variance and low conservation of ‘connectivity’ (Fig. 3A-C), and integumental patterns can vary greatly, even amongst closely related species.^54–56^ Our comparative gene co-expression analyses in single cells can therefore provide important clues whether a certain patterning process is likely to be dominated by changes in the extracellular environment, or if cell-intrinsic factors are also important to consider for its amenability to evolutionary change.

### 3.3 Cell types and cell states, in development and evolution

Lastly, looking at our module-based clustering of pseudobulks, we often observed discrepancies in cluster composition, compared to our original cell type assignments. This holds especially true for pseudobulks of lateral plate mesoderm origin (Fig. 3D, Fig. 4A, B). There, many of our prior assignments – based on differential expression analysis and marker gene identification – no longer seem to concur with the transcriptional clustering of our gene co-expression modules. As a result, pseudobulks of different assigned cellular identities, e.g. chondrocytes, mesenchyme or interdigit, start to intermingle. Upon closer inspection, this trend seems to be driven – to a large extent – by the differential activities of modules ‘green’ and ‘midnightblue’, i.e. ‘cell cycle’ and ‘cellular respiration’, respectively (Fig. 3D). Both of these modules clearly seem more indicative of cell state, than cell type.^18^ Therefore, using co-expression module detection on single-cell data appears to reveal commonalities in the expression dynamics of groups of genes that otherwise might go unnoticed. For example, if relying on differential expression analyses alone, i.e. by contrasting each of the populations against the rest of the cells, groups of genes with broad expression patterns will most likely not be detected as markers of a given cell population.^57^ The issue seems particularly relevant for genes that relate to cell state, rather than cell type, as module-based cell state signatures of gene expression can be shared by a variety of different cell types (Fig. 3D, Fig. 4A, B).

Accordingly, we advocate for a multilayered approach when assigning cellular identifiers to scRNA-seq data, where a combination of differential expression analyses, cluster-independent gene coexpression module detection, and prior knowledge of the biological system at hand is taken into consideration. At a broader scale, even in samples from embryonic stages, the data will generally have the tendency to sort according to developmental lineage, cell type, and only then cell state. The last two categories especially, however, can be difficult to disentangle during development. Many cell types can often be present in multiple stages of differentiation, with rare trajectional intermediates – or transitional stages – interspersed in between.^58–61^ Whether those themselves should be considered distinct cell types, or rather cell states, can be a matter of debate.^10,18,62^ Cleary, though, accounting for more general, lineage-independent cell states should result in a more comprehensive appreciation of the respective cell type behaviors, with e.g. ‘cell cycle’ expected to be a dominant signature in any growing tissue. This will only become more relevant, as scRNA-seq studies continue to expand into investigating the impacts of different genetic backgrounds, or environmental variables.^63–65^

Overall, we observe that our comparative gene co-expression module approach represents a valuable addition to discriminate distinct cell states, some of which can be shared amongst different cell types or even distinct developmental lineages. Especially among early, undifferentiated tissues, these module signatures can contain important temporal information across samples, but also – for more mature cell types – signals relevant for comparisons between distantly related species, mutant backgrounds, and environmental parameters.

## 4 EXPERIMENTAL PROCEDURES

### 4.1 Sampling and Data Sources

We sampled complete forelimbs at stages HH21, HH24 and HH27. Tissue dissociation and *10x* Genomics Chromium 3’ Kit library preparation was performed as reported previously.^21^ We obtained for HH21 / HH24 / HH27 a total of 2990 / 6345 / 2189 cells, with median UMI counts of 2365 / 2215 / 1315 and median number of genes detected of 978 / 952 / 637 per cell. Raw sequencing data and UMI count matrices are available under GEO accession *‘in-process’*.

Publicly available datasets used in this study were mouse E9.5 and E10.5 (GEO accession: GSE149368)^20^; mouse E11.5, E13.5, E15.5 and E18.5 (GEO accession: GSE142425,)^19^; and chicken HH25, HH27 and HH29 (GEO accession: GSE130439).^21^

### 4.2 3’UTR elongation and improved chick genome annotation

To elongate 3’UTR annotations, we used stage HH11, HH14, HH21/22, HH25/26, HH32 and HH36 whole embryo bulk RNA-seq datasets.^23^ RNA-seq reads were processed and mapped individually for each stage, filtered and down-sampled to 40 million pairs of mapped reads per sample. Resulting BAM files were merged and used to generate transcript models with Cufflinks.^66^ The newly calculated transcript models were then processed for 3’ UTR elongation.

Elongation of existing GRCg6a 3’ UTR annotations was conducted with the following logic: We only considered transcript models with expression >1 FPKM, which overlapped only one original gene annotation track, and where the original 3’ UTR annotation was shorter than the novel model. 3’ UTR elongation was capped at a maximum of 5,000 bp, and was shortened accordingly, if it resulted in any overlap with a neighboring gene. A total of 3132 3’UTRs were elongated in such way. Additionally, we realized that with the migration form Gallus_gallus-5.0, 225 gene stable IDs associated with a gene name were now absent from GRCg6a. Using a combination of BLAST^67^ and the GenomicRanges and IRanges packages^68^ in R, we managed to recover 62 of these genes and appended them to our modified GRCg6a annotation.

### 4.3 Single-cell data pre-processing

All chicken samples were processed with *CellRanger (10x* Genomics), using our improved GRCg6a genome annotation. Chicken and mouse UMI count matrices were processed, with cells filtered for quality based on total and relative UMI counts and percentage of mitochondrial UMIs. UMI matrices for E9.5 and E10.5 samples are already filtered for total UMI counts and mitochondrial counts. Due to the overall size of these two data sets, we randomly subsampled 25% of the single cell transcriptomes, to have datasets of comparable sizes. Moreover, we excluded 4412 cells from the first replicate of the E9.5 sample showing abnormal haemoglobin genes expression.

### 4.4 Data normalization and correction

UMI count data was normalized cell-wise using Seurat v3.1.4^25^ with a scale factor of 10000 and then log-transformed. Total UMI count, proportion of mitochondrial UMIs and cell cycle stage scores^21,69,70^ were then used as variables to regress using the function “SCTransform” from Seurat. Moreover, for samples E9.5 and HH29 sequencing batch effects were also regressed. It is important to note that cell cycle correction is only applied to calculate PCs, tSNEs and clusters, but not for differential expression analyses and all other analyses.

### 4.5 Dimensionality reduction, cell clustering and cluster annotation

We performed principal component analysis (PCA) using Seurat’s “RunPCA” with default options. Significant PCs were determined for each sample as those falling outside of a Marchenko-Pastur distribution^71^ and tSNEs were produced to retain and represent the global structure of the data.^72, 73^ To infer cell clusters, we identified the nearest neighbors of each cell, using the first significant PCs. We calculated a hierarchical tree of clusters using “BuildClusterTree” based on significant PCs and identified ‘sister tips’ and performed differential expression tests on each of them. If two clusters showed less than 5 genes differentially expressed, they were merged, and the process repeated with a new tree of clusters. Differential expression analyses were performed with the MAST^74^ implementation in Seurat. Using “FindVariableFeatures”, we selected highly variable genes with a standardized variance larger than the sample median. For making comparisons across clusters we used normalized but “uncorrected” data, using the δ(S-G2M) as a latent variable. We only tested highly variable genes expressed in at least 25% of the cells in either cell population. Only genes with an adjusted p-value < 0.05 and log2 fold change > 0.5 were considered as differentially expressed. Differentially expressed genes were then used as “marker genes” for cell cluster annotation, in combination with spatial gene expression data repositories like *Geisha* (Chicken Embryo Gene Expression Database)^75^ and *MGI* (Mouse Gene Expression Database),^76^ as well as GO-term enrichment analyses.^77^

Data integration into a single tSNE per species was conducted using transformed data and “IntegrateData” with its related functions. We used as anchors all the shared expressed genes for the mouse and 3000 highly variable genes for the chicken, with 20 dimensions and a k.filter of 100. PCA and tSNE were calculated as above.

### 4.6 *R* package “scWGCNA”

The main analytical workflow presented in this paper is contained within a newly developed *R* package, “scWGCNA”, and is available on GitHub with accompanying documentation at https://github.com/CFeregrino/scWGCNA It consists of three main functions, as outlined below.

**‘Pseudocell’ function** - To increase robustness, we define so-called ‘pseudocells’. The 10 nearest neighbors (NN) of each cell were calculated in the PCA space using “FindNeighbors”. From each of the previously calculated cell clusters, 20% of the cells were chosen randomly as seed cells. In order to maximize the number of cells aggregated into pseudocells, we perform a sampling of 50 sets of randomly chosen seed cells and choose the set with the largest NN count. Moreover, seed cells typically share some of their NN with other seed cells, for which we do an iterative cell distribution step. First, to avoid “greedy” seed cells, starting with the seed cell with the lowest amount of remaining NN, one of its NN is chosen at random. The chosen cell is removed from the universe of cells and its assigned seed cell is recorded. Once all cells have been distributed to a seed cell, we use the function “AverageExpression” to aggregate the scaled expression of each resulting pseudocell.

**‘Iterative WGCNA’ function** - We calculate highly variable genes from normalized single cell data using “FindVariableFeatures” and the “mvp” method with cutoffs of minimal 0.25 dispersion and minimal 0 expression. Then, with pseudocell expression data, a soft thresholding power is selected to calculate an adjacency matrix using “pickSoftThreshold” in WGCNA with the bidweight midcorrelation method and a signed network type.^24^ The WGCNA analyses itself occurs in a recursive manner. A topological overlap matrix is produced from pseudocell expression data with the function “TOMsimilarityFromExpr”, with previously calculated soft thresholding power and bidweight midcorrelation. A hierarchical clustering tree is then computed using the topological overlap distances. A series of cut heights are set in steps of 0.0001 around (+- 0.0005) of a height of 99% of the range between the 5th percentile and the maximum heights on the clustering tree. The size of the detected modules for each cut height is recorded, and the height producing the smallest – or no – gray module (i.e. unassigned genes), and the same number of modules as the previous iteration (or 20, in the first run) is selected. Once a height is selected, modules are detected, and module membership of each gene is calculated using “geneModuleMembership”. Genes not assigned, or without significant module membership, are removed, and the remaining ones are used to start the process again. Once all remaining genes have significant module membership, eigengenes and average expression of each module are calculated in single-cell space, and GO-term enrichment analyses for each module us performed using Limma.^21, 77^ All output is contained within a single HTML file, with averaged module expression plotted on tSNEs. Networks are visualized using R packages ‘network’^78^ and ‘GGally’,^79^ with edge thicknesses and intensities scaled module-wise to represent topological overlap.

**‘Comparative WGCNA’ function** - We use pseudocell data of both reference and test datasets. We subset the modules to contain only genes present as high confidence 1-to-1 orthologues, using orthologous genes list from ENSEMBL BioMart.^80^ Using the “goodGenes” function from WGCNA, we filter genes based on expression and variance in all test samples. Conservation test is performed by “modulePreservation”, with filtered module assignments, bidweight midcorrelation, a maximal gold modules size of 300, and 20 permutations.^31^ The overall conservation Zsummary and median rank, as well as the density and connectivity conservation Zsummary are summarized in a single HTML file.

## ACKNOWLEDGMENTS

The authors wish to thank Henrik Kaessmann and his lab for hosting CF for his EMBO STF, Virginie Ricci and Fabio Sacher for beta-testing the “scWGCNA” *R* package, Lila Allou and Stefan Mundlos for making mouse scRNA-seq data available prior to publication, Christian Beisel and the joint Genomics Facility Basel for help with single-cell sequencing, and all members of our group for useful discussions. All calculations were performed at sciCORE (http://scicore.unibas.ch/), scientific computing center at the University of Basel. This work was supported by an EMBO short-term fellowship to CF (Fellowship Number: 8593), and research funds from the Swiss 3R Competence Centre (3RCC grant OC-2018-005), the Swiss National Science Foundation (SNSF project grant 310030_189242), and the University of Basel to PT.

## AUTHOR CONTRIBUTIONS

CF and PT designed the study, CF performed all wet-lab work and data analyses and wrote the “scWGCNA” *R* package, CF and PT wrote the paper.

## Supplemental Figure 1

**Supplemental Figure 1.**
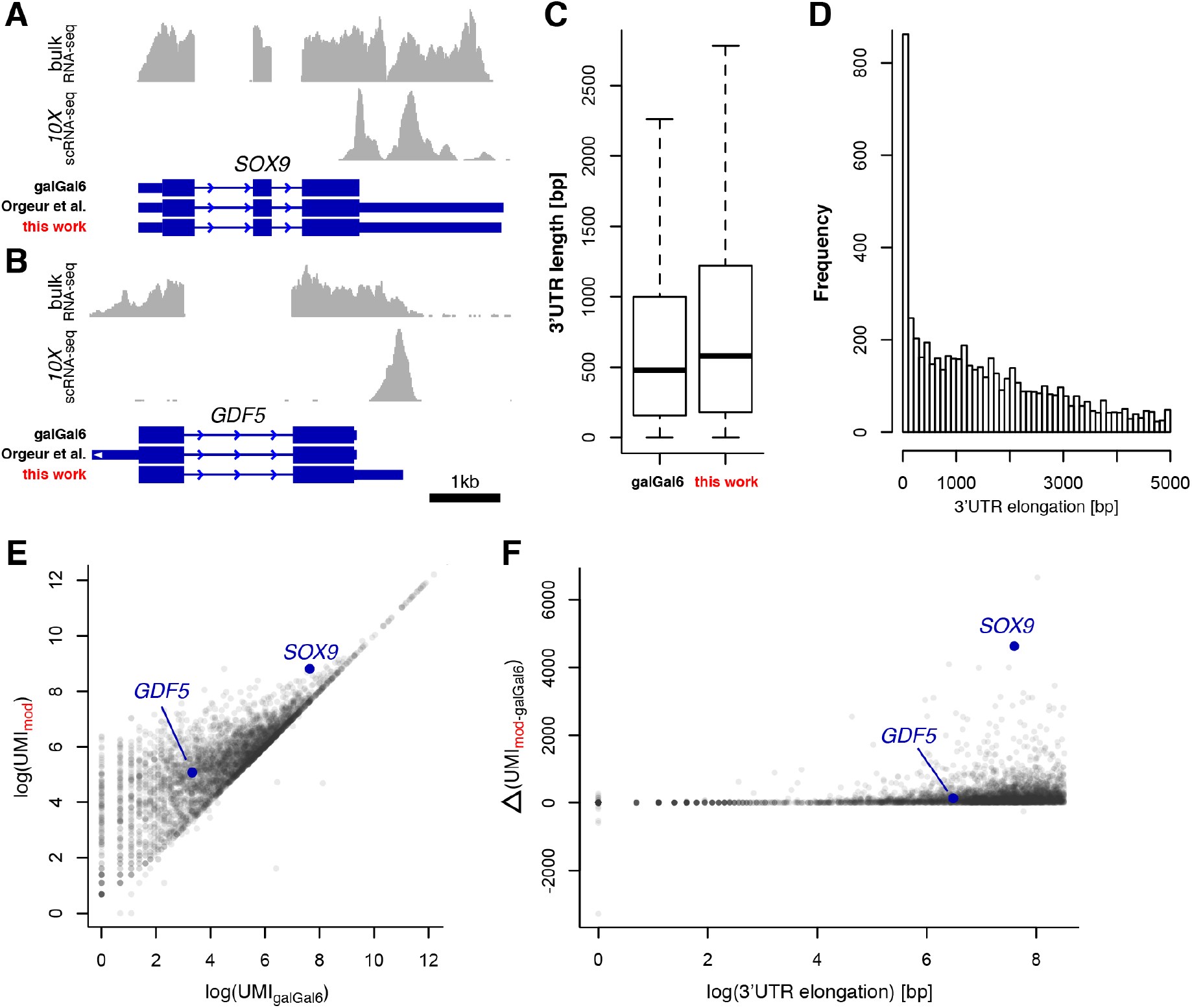
Elongating 3’UTR annotations in the chicken genome. (A, B) Genome browser visualizations of mapped reads, stemming from either bulk (top) or *10x* Genomics single-cell (bottom) RNA-seq experiments, at the *SOX9* (A) and *GDF5* (B) locus. The respective gene models are displayed from the original GRCg6a annotation (galGal6), the annotation of Orgeur and colleagues^81^ and the annotation produced in this work. (C) Boxplots of 3’UTR lengths of the original GRCg6a and our annotation. (D) Histogram showing the distribution of elongation lengths for all elongated 3’UTRs. (E) Resulting changes in UMI counts, when mapping our HH31 sample to either the original GRCg6a or our modified annotation. For the large majority of genes, UMI counts either stay unchanged or show an increase with our annotation. We visually inspected the few genes that showed a reduced UMI count and realized that these UMIs had been attributed to another gene or transcript model in our second *CellRanger* run, i.e. they were not lost due to misguided 3’UTR elongations. (F) Correlation of 3’UTR elongation length and increase in UMI counts. Only a weak effect was found of the amount of 3’UTR lengthening on the gained UMI counts.

**Supplement 1**

HTML output of the ‘Iterative WGCNA’ function

**Supplement 2**

HTML output of the ‘Comparative WGCNA’ function

