## Supplement 1 for "Assessing evolutionary and developmental transcriptome dynamics in homologous cell types"

WGCNA workflow, E15.5

Code 

- Show All Code
- Hide All Code
- Download Rmd

### WGCNA workflow, E15.5

###### Christian Feregrino

Date: 03.02.21

We use WGCNA to run an iterative analysis on a data set. We are calculating HVG in the script, but using a threshold of 0.25 for the dispersion.

#### Pre-analysis

We need to first set up our working environment.

```
# WGCNA
#BiocManager::install("WGCNA")
library("WGCNA")
library("Seurat")
library("ggplot2")
library("gridExtra")
library("GGally")
library("network")

# The following setting is important, do not omit
options(stringsAsFactors = FALSE);
```

Get our single cells and the pseudocells to be able to run the analysis

```
# placeholder for the variables and options we will get from an upper level pipeline

setwd("~/projects/scMmGg_Oct20/WGCNA/")

my.date = "091120"
project_name = "E15"

p.Wdata = readRDS(file = "./../robjects/E15_seurat_271020.rds")

gnames = readRDS("./../data/mouse_gnames.rds")

Wdata = readRDS(file = paste0("./data/",project_name,"_pseudocells_",my.date,".rds"))

# The subset in form of cell identities, F if using the whole sample
my.subset = F
# Variable genes, or genes to use, F if the genes will be calculated in the script
my.vargenes = F

my.assay = "SCT"

my.dr = "tsne"

dir.create("./WGCNA_results", showWarnings = F)

my.sp = "Mm"
```

We need the data in either seurat or data-frame format. This must contain at least 20 samples (no problems with sc data), according to the documentation of the package itself.

```
# If we need to subset the data
if (my.subset) {
  p.Wdata = subset(p.Wdata, idents=my.subset)
}

# If no variable genes are provided
nonex = which(apply(p.Wdata@assays$RNA@counts, 1, function(x) length(which(x >0))) < 10)

if (my.vargenes == F) {
  # First get rid of non-expressed genes
  p.Wdata = subset(p.Wdata, features = rownames(p.Wdata@assays$RNA)[-nonex])
  
  # Find the variable genes
  p.Wdata=FindVariableFeatures(p.Wdata, dispersion.cutoff = c(0.25,Inf), mean.cutoff = c(0,Inf), selection.method = "mvp", assay = "RNA")
  VariableFeaturePlot(p.Wdata, assay = "RNA")
  Expr = VariableFeatures(object = p.Wdata, assay = "RNA")
  
} else { Expr = my.vargenes }
```

```
Calculating gene means
0%   10   20   30   40   50   60   70   80   90   100%
[----|----|----|----|----|----|----|----|----|----|
**************************************************|
Calculating gene variance to mean ratios
0%   10   20   30   40   50   60   70   80   90   100%
[----|----|----|----|----|----|----|----|----|----|
**************************************************|
```

```
# if (length(my.ortho) > 1) {
#   Expr = Expr[which(Expr %in% my.ortho[,1])]
#   Expr = Expr[ which(Expr %in% rownames(p.Wdata@assays$RNA)[-nonex]) ]
# }
  
datExpr=Wdata@assays$RNA@counts[Expr,]

# Check the length
print(paste0("We have ", length(Expr), " genes in the variable genes object"))
```

```
[1] "We have 2967 genes in the variable genes object"
```

```
# Check the size and transform
dim(datExpr)
```

```
[1] 2967  513
```

```
datExpr = t(as.matrix(datExpr))
```

Now we need to calculate the soft threshold power. First it calculates the similarity and then transforms this similarity to a weighted network. The scale-free topology is calculated for each of the powers.

We choose the smallest power for which the scale-free topology fit index reaches 0.90. If none of the powers reaches 0.90, we take the one with the maximum, as long as we have a number above 0.75. If none of them reaches at least 0.75 we need to check our dataset.

```
# Choose a set of soft-thresholding powers
powers = c(c(1:10), seq(from = 12, to=30, by=2))

# Call the network topology analysis function
sft = pickSoftThreshold(datExpr, powerVector = powers, verbose = 0, networkType = "signed", corFnc = "bicor")
```

```
bicor: zero MAD in variable 'x'. Pearson correlation was used for individual columns with zero (or missing) MAD.bicor: zero MAD in variable 'y'. Pearson correlation was used for individual columns with zero (or missing) MAD.
```

```
# Plot of the scale-free topology fit index as a function of the soft-thresholding power
plot(sft$fitIndices[,1], -sign(sft$fitIndices[,3])*sft$fitIndices[,2],
     xlab="Soft Threshold (power)",ylab="Scale Free Topology Model Fit,signed R^2",type="n",
     main = paste("Scale independence"))
text(sft$fitIndices[,1], -sign(sft$fitIndices[,3])*sft$fitIndices[,2],
     labels=powers,col="red")
```

```
which((-sign(sft$fitIndices[,3])*sft$fitIndices[,2]) > 0.9)
```

```
 [1]  5  6  7  8  9 10 11 12 13 14 15 16 17 18 19 20
```

```
# this line corresponds to using an R^2 cut-off of h
abline(h=0.85,col="red")
```

```
# These are the scale-free topology indexes
indexes = (-sign(sft$fitIndices[,3])*sft$fitIndices[,2])

# If we don't have aany index above 0.75, we stop the script
if ( !any( indexes > 0.75 ) ) {
  print("The scale-free topology index didn't reach 0.75 with any of the chosen powers, please consider changing the samples")
  # quit(save = "no", 1, F)
}

# Take the smalles power that gives us an index over 0.9, or the highest index if we don't reach 0.9
if ( any( indexes > 0.9 ) ) {
    my.power = sft$fitIndices$Power[min(which(indexes > 0.9))]
} else { my.power = sft$fitIndices$Power[which.max(indexes)] }

print(paste0("Or power is ", my.power))
```

```
[1] "Or power is 5"
```

#### WGCNA analysis

Now, the following piece of code will run WGCNA iteratively, to end up with out final modules. The iterations follow these steps:

- Tree cutting for module calculation
  - Calculate an adjacency matrx from the data, then turn it into topological overlap and then into a distance matrix
  - Calculate the tree based on the topological overlap distance
  - Calculate the automatic height to cut out the 0.05 quantile
  - Generate a matrix where we calculate the amount of modules, size of de modules and wheter if we have a grey module based on:
    - Different minimum module sizes, arbitrarely set to 7:30
    - Different cut-heiughts going 0.0005 up and down from the automatic height in steps of 0.0001
  - Check if any combination of the parameters will get rid of the grey module
    - If we only get grey modules, subset the matrix for the height at which the grey module is the smallest
    - If we have a combination without grey module AND we have at least the same number of modules as in the beginning, we subset for whichever those heights are
  - Take wichever min module sizes gives us at least the same amount of clusters as in the beginning, if none, then the highest
  - Chose the max of the reminding min module sizes.
- Calculate the actual modules
- Calculate the eigengenes
- Calculate the module membership per gene
- Calculate the p.value of the membership to a given module
- Get rid of the genes in the grey module
- Delete the genes that are not significantely associated with their module
- Save the remaining geneset
- Print how many genes were deleted due to significance
- Unless 0 genes were deleted in the last step, update the expression matrices and beginn again.

ONLY THE FIRST TWO ITERATIONS ARE DIFFERENT.

FIRST:

- During the tree cutting
  - Set the minimum module size to arbitrary 15
  - Subset for the heights that get rid of at least 50% of the genes with that min module size
  - Choose the height that gives us the most clusters

SECOND:

- Set the resulting number of modules as the ground number of modules

```
# Running WGCNA iteratively

my.Clnumber = 20
change = 0
genesets=list()
nonsig = 1

while(nonsig != 0) {
  
  # Turn adjacency into topological overlap (high overlap if they share the same "neighborhood")
  TOM=TOMsimilarityFromExpr(datExpr,networkType = "signed", TOMType = "signed", power = my.power, corType = "bicor")
  
  #Put the names in the tree
  colnames(TOM) <- gnames[colnames(datExpr),"Name"]
  
  rownames(TOM) <- gnames[colnames(datExpr),"Name"]
  
  #Make it a distance
  dissTOM = 1-TOM
  
  # Call the hierarchical clustering function
  geneTree = hclust(as.dist(dissTOM), method = "average")
  
  
  # Here I calculate the cutting height. Using the same formula and approach that the WGCNA package uses for the automatic function
  nMerge = length(geneTree$height) # The whole height of the tree
  refQuantile = 0.05 # What's the quantile that we want to exclude
  refMerge = round(nMerge * refQuantile) 
  refHeight = geneTree$height[refMerge]
  cutheight = signif(0.99 * (max(geneTree$height) - refHeight) + refHeight,4)
  
  # We construct THE TABLE that will help us make decisions
  # Min cluster sizes, from 7 to 30
  x=seq(7,30,1)
  # The height, up and down from the calculated height. We expect some "No module detected"
  y=seq(cutheight-0.0005,cutheight + 0.0005,0.0001)
  
  # The actual dataframe
  w=data.frame()
  # Populate, with i=min cluster size, j=cutting height, z=total number of clusters, z.1.=what's the first cluster? 0 is grey 1 is something else,
  # z.1.'=what's the size of the first cluster?
  for (i in x) {
    for (j in y) {
      sink("aux")
      z=table(cutreeDynamic(dendro = geneTree,  method="tree", minClusterSize = i, deepSplit = T, cutHeight = j, verbose = 0))
      sink(NULL)
      v=data.frame(i,j,dim(z),names(z[1]),unname(z[1]))
      w=rbind(w,v)
    }
  }
  
  # The height is then the one where we have the least number of genes in the first cluster
  my.height = w$j[which(w$unname.z.1..==min(w$unname.z.1..))]
  
  # Since different heights can give us the minimum grey size, we chose the computed height, if present, or the highest one.
  if (cutheight %in% my.height) {
    my.Clsize = w[which(w$j == cutheight),]
  } else { my.Clsize = w[which(w$j == max(my.height)),] }
  
  
  # This is to know, if we're looking for a minimum of cluster numbers
  # If we still have a lot of genes, we don't want to limit the number of clusters
  # if ( ((dim(datExpr)[2]) / length(Expr)) > 0.6 ) {change = 0}
  
  # If this is the first iteration after 0.6 of the genes are gone, we assign the number of clusters (and an extra for the grey in the case)
  if (change == 2) {
    my.Clnumber = length(table(dynamicColors)) + 1
  }
  # Count another iteration
  change = change + 1
  
  # If we don't have a gray cluster anymore, then we subset for those rows, and set a new height. ONLY if we get the same amount of clusters!
  if (any(w$names.z.1.. == 1)) { #any combination gives us no grey
    
    my.Clsize = w[which(w$names.z.1.. == 1),] # Take all combinations that gives us no grey
    
    if (any(my.Clsize$dim.z. >= (my.Clnumber -1) )) { # If there is any giving us the determined amount or more
      my.Clsize = my.Clsize[which(my.Clsize$dim.z. >= (my.Clnumber - 1) ),,drop=F] # Subset for those
    } else { my.Clsize = my.Clsize[which(my.Clsize$dim.z. == max(my.Clsize$dim.z.)),,drop=F] } # Or for the highest
    
    # Take the ones with the smallest number of clusters
    my.Clsize = my.Clsize[which(my.Clsize$dim.z. == min(my.Clsize$dim.z.)),,drop=F]
    # Take the one with the highest min cluster size
    my.Clsize = my.Clsize[which(my.Clsize$i == max(my.Clsize$i)),,drop=F]
    
    if (cutheight %in% my.Clsize$j) { # if original computed height is in,
      my.height = cutheight # take it
    } else { my.height = max(my.Clsize$j) } # Otherwise, the highest height
    
    my.Clsize = max(my.Clsize$i)
    
  }
  
    # Subset the table again, for those sizes that will gives the same number of clusters or more. IF NONE, use the highest number
  if (!any(w$names.z.1.. == 1)){
    if (any(my.Clsize$dim.z. >= my.Clnumber)) {
      my.Clsize = my.Clsize[which(my.Clsize$dim.z. >= my.Clnumber),,drop=F]
    } else {
      my.Clsize = my.Clsize[which(my.Clsize$dim.z. == max(my.Clsize$dim.z.)),,drop=F]}
    
   # Take the ones with the smallest number of clusters
    my.Clsize = my.Clsize[which(my.Clsize$dim.z. == min(my.Clsize$dim.z.)),,drop=F]
    # Take the one with the highest min cluster size
    my.Clsize = my.Clsize[which(my.Clsize$i == max(my.Clsize$i)),,drop=F]
    
    if (cutheight %in% my.Clsize$j) { # if original computed height is in,
      my.height = cutheight # take it
    } else { my.height = max(my.Clsize$j) } # Otherwise, the highest height
    
    my.Clsize = max(my.Clsize$i)
  }
  
  # If we still have more than 60% of the genes, we just use the min size of 15 regardles
  if ( change < 3 ) {
    my.Clsize = 15
    my.height = cutheight
  }
  
  dynamicMods = cutreeDynamic(dendro = geneTree,  method="tree", minClusterSize = my.Clsize, deepSplit = T, cutHeight = my.height)
  
  #dynamicMods = cutreeDynamic(dendro = geneTree, distM = dissTOM, deepSplit = 2, minClusterSize = minModuleSize)
  
  table(dynamicMods)
  
  # Convert numeric lables into colors
  dynamicColors = labels2colors(dynamicMods)
  table(dynamicColors)
  
  # Plot the dendrogram and colors underneath
  plotDendroAndColors(geneTree, dynamicColors, "Modules",
                      dendroLabels = NULL,
                      cex.dendroLabels = 0.6,
                      addGuide = TRUE,
                      main = "Gene dendrogram and module colors",
                      guideAll = F)
  
  par(mfrow=c(1,1))
  
  # Calculate eigengenes
  MEList = moduleEigengenes(as.matrix(datExpr), colors = dynamicColors)
  
  MEs = MEList$eigengenes
  
  # Calculate the module membership
  geneModuleMembership = as.data.frame(signedKME(datExpr, MEs))
  MMPvalue = as.data.frame(corPvalueStudent(as.matrix(geneModuleMembership), nrow(datExpr)))
  
  # We're gonna make a list, where we keep the genes that are significantly associated with each module
  x=c()
  xy=list()
  
  # We also need a vector with all the dynamic colors
  dcols = 1:length(levels(as.factor(dynamicColors)))
  
  # Getting rid of the grey module
  grey.genes = length(which(dynamicColors == "grey"))
  if (any(levels(as.factor(dynamicColors)) == "grey")) {
    dcols = dcols[-which(levels(as.factor(dynamicColors)) == "grey")]
  }
  
  # Run the loop to get the genes
  for (i in dcols) {
    modGenes = rownames(MMPvalue)[which(dynamicColors==levels(as.factor(dynamicColors))[i] & MMPvalue[,i]<0.01)]
    x=c(x,modGenes)
    xy[[i]]=modGenes
    #print(paste0(levels(as.factor(dynamicColors))[i]," ",length(modGenes),
    #" of ", length(which(dynamicColors==levels(as.factor(dynamicColors))[i]))))
    #print(gnames[modGenes,2])
  }
  
  # Make a new list, where we keep ALL the gens thar are left from the iteration, that will be used to make the new object. To keep track
  genesets[[length(genesets)+1]] = colnames(datExpr)
  
  # Give me a message saying how many genes are gone this time
  cat( paste0( grey.genes, " genes not assigned to any module.", '\n',
                 length(which(!(colnames(datExpr)%in%x))) - grey.genes, " genes excluded due to significance."))
  # Save this also, cause if it's 0 then we stop the whole thing
  nonsig = length(which(!(colnames(datExpr)%in%x)))
  
  # If it ain't 0, subset the dynamic colors and the expression data
  if (length(which(!(colnames(datExpr)%in%x))) != 0) {
    dynamicColors=dynamicColors[-which(!(colnames(datExpr)%in%x))]
    datExpr=datExpr[,-(which(!(colnames(datExpr)%in%x)))]
  } 

}
```

```
TOM calculation: adjacency..
..will not use multithreading.
 Fraction of slow calculations: 0.000000
..connectivity..
..matrix multiplication (system BLAS)..
..normalization..
..done.
0 genes not assigned to any module.
528 genes excluded due to significance.TOM calculation: adjacency..
..will not use multithreading.
 Fraction of slow calculations: 0.000000
..connectivity..
..matrix multiplication (system BLAS)..
..normalization..
..done.
```

```
0 genes not assigned to any module.
135 genes excluded due to significance.TOM calculation: adjacency..
..will not use multithreading.
 Fraction of slow calculations: 0.000000
..connectivity..
..matrix multiplication (system BLAS)..
..normalization..
..done.
```

```
0 genes not assigned to any module.
161 genes excluded due to significance.TOM calculation: adjacency..
..will not use multithreading.
 Fraction of slow calculations: 0.000000
..connectivity..
..matrix multiplication (system BLAS)..
..normalization..
..done.
```

```
0 genes not assigned to any module.
37 genes excluded due to significance.TOM calculation: adjacency..
..will not use multithreading.
 Fraction of slow calculations: 0.000000
..connectivity..
..matrix multiplication (system BLAS)..
..normalization..
..done.
```

```
0 genes not assigned to any module.
36 genes excluded due to significance.TOM calculation: adjacency..
..will not use multithreading.
 Fraction of slow calculations: 0.000000
..connectivity..
..matrix multiplication (system BLAS)..
..normalization..
..done.
```

```
0 genes not assigned to any module.
9 genes excluded due to significance.TOM calculation: adjacency..
..will not use multithreading.
 Fraction of slow calculations: 0.000000
..connectivity..
..matrix multiplication (system BLAS)..
..normalization..
..done.
```

```
0 genes not assigned to any module.
8 genes excluded due to significance.TOM calculation: adjacency..
..will not use multithreading.
 Fraction of slow calculations: 0.000000
..connectivity..
..matrix multiplication (system BLAS)..
..normalization..
..done.
```

```
0 genes not assigned to any module.
3 genes excluded due to significance.TOM calculation: adjacency..
..will not use multithreading.
 Fraction of slow calculations: 0.000000
..connectivity..
..matrix multiplication (system BLAS)..
..normalization..
..done.
```

```
0 genes not assigned to any module.
2 genes excluded due to significance.TOM calculation: adjacency..
..will not use multithreading.
 Fraction of slow calculations: 0.000000
..connectivity..
..matrix multiplication (system BLAS)..
..normalization..
..done.
```

```
0 genes not assigned to any module.
1 genes excluded due to significance.TOM calculation: adjacency..
..will not use multithreading.
 Fraction of slow calculations: 0.000000
..connectivity..
..matrix multiplication (system BLAS)..
..normalization..
..done.
```

```
0 genes not assigned to any module.
0 genes excluded due to significance.
```

```
p.MEList = MEList

raw.datExpr = p.Wdata@assays$RNA@scale.data[colnames(datExpr),]

raw.datExpr = t(as.matrix(raw.datExpr))

raw.MEList = moduleEigengenes(raw.datExpr, colors = dynamicColors)
  
p.MEList = raw.MEList
```

#### Modules of co-expression

For single cells We can see what are the expression levels of our co-expression modules. If we provided a seurat object. We look, in this case, at a tSNE

```
xx=list()
yy=c(levels(as.factor(dynamicColors)))
for (i in 1:length(yy)) {
  toplot = data.frame(p.Wdata@reductions[[my.dr]]@cell.embeddings)
  
  xx[[i]] = ggplot(toplot[order(p.MEList$averageExpr[,i]),],
                   aes_string(x=colnames(toplot)[1], y=colnames(toplot)[2])) +
  geom_point(aes_string(color=p.MEList$averageExpr[order(p.MEList$averageExpr[,i]),i]), size=2) +
  scale_size(range = c(1, 1)) +
  theme_void() + theme(legend.position="none") +
  scale_colour_gradientn(colours = c("gray90", "gray90", yy[i], yy[i])) +
  labs(colour=levels(as.factor(dynamicColors))[i])
}
grid.arrange(grobs=xx, ncol=4)
```

Create a list object with all the essential data from the analyses

```
WGCNA_data = list()
WGCNA_data[["datExpr"]] = datExpr
WGCNA_data[["dynamicMods"]] = dynamicMods
WGCNA_data[["MEList"]] = MEList
WGCNA_data[["MEs"]] = MEs
WGCNA_data[["modGenes"]] = xy
WGCNA_data[["modPlots"]] = xx
WGCNA_data[["genesets"]]= genesets
WGCNA_data[["TOM"]]= TOM
WGCNA_data[["adjacency"]]= adjacency

saveRDS(WGCNA_data, file = paste0("./data/",project_name,"_WGCNA_data.rds"))
```

To create the visualizations of the actual co-expression networks, we use this code. It also creates files that can be read into cytoscape

```
# we create a new directory where the network files will be
my.dirname = paste0("./data/",project_name,"WGCNA_networks/")

dir.create(my.dirname)
```

```
'./data/E15WGCNA_networks' already exists
```

```
modules=levels(as.factor(dynamicColors)) # The modules

# To generate the files necessary for the network visualization in cytoscape
for (i in 1:length(modules)) {
  mod=is.finite(match(dynamicColors, modules[i])) # Each module
  cyt = exportNetworkToCytoscape(TOM[mod,mod], 
                                 edgeFile = paste( paste0(my.dirname,"CytoscapeInput-edges-"),
                                                  paste(modules[i], collapse="-"), ".txt", sep=""), # The name of the edge file
                                 nodeFile = paste( paste0(my.dirname,"CytoscapeInput-nodes-"),
                                                  paste(modules[i], collapse="-"), ".txt", sep=""), # The node file
                                 weighted = TRUE, #Puts the weights
                                 threshold = 0.0, #Threshold for adjacency, 0=all genes
                                 nodeNames = colnames(datExpr)[mod], #The names of the nodes/genes
                                 altNodeNames = gnames[colnames(datExpr)[mod],2], #Other names
                                 nodeAttr = geneModuleMembership[colnames(datExpr)[mod],i]) #Some more info about the nodes
}

# We build a list where we keep the network plots
netplots=list()

lcols = c(rep("black", length(unique(dynamicColors))))
lcols[(apply(col2rgb(levels(as.factor(dynamicColors))), 2,
               function(x) (x[1]*0.299 + x[2]*0.587 + x[3]*0.114)) < 75)] = "white"

mynets = list()

# A loop that makes the network plots in R
for (i in 1:length(xx)) {
  # Load directly edges tables from the file we created
  mynetwork = read.table(paste0(my.dirname,"CytoscapeInput-edges-",levels(as.factor(dynamicColors))[i],".txt"),
                       header = T, stringsAsFactors = F, fill=T)
  # We get rid of all the edges that have a very small weight. The cutoff set to keep at least ONE edge on the nodes
  x = max( c( min(aggregate(mynetwork$weight, by = list(mynetwork$fromNode), max)$x),
              min(aggregate(mynetwork$weight, by = list(mynetwork$toNode), max)$x) ) )
  mynetwork=mynetwork[-which(mynetwork$weight < x),]
  
  # We rescale the weights so that we have them from 0 to 1
  mynetwork$weight01 = rescale01(mynetwork[,3])
  # And then multiply them by 2, to give the heavy edges a 2 thickness, we ad 0.2 to not have completelly invisible edges
  mynetwork[,3] = (rescale01(mynetwork[,3]) * 2) + 0.2
  # Convert this edgelist into a network object
  mynet = network(mynetwork, matrix.type="edgelist", ignore.eval=F)
  
  # Load in the nodes table
  mynodes = read.table(paste0(my.dirname,"CytoscapeInput-nodes-",levels(as.factor(dynamicColors))[i],".txt"),
                     header = T, stringsAsFactors = F, fill = T)
  rownames(mynodes) <- mynodes$nodeName
  # put in the membership value, from the ModuleMembership table, and rescale that from 0 to 1.
  mynodes$membership = rescale01(geneModuleMembership[rownames(mynodes),i]) 
  # We multiply by 30 to give a good range of sizes, plus 1 to avoid innexistant nodes
  mynodes$membership= (mynodes$membership*30)+1
  
  mynet = set.vertex.attribute(mynet, "membership", mynodes[network.vertex.names(mynet),"membership"])
  
  mynets[[i]] = mynet
  
}
```

To check the results module by module, we report some GO terms and individual plots, with module sizes and gene names. First the GO terms analyses

```
# We need these two packages to run GO analyses, on the chicken. Replace for any other GO functions
library(paste0("org.",my.sp,".eg.db"),character.only = T)

library(limma)

#Get the ENSEMBL to ENTREZ list from the BioMart

my.ENS2EG = get(paste0("org.",my.sp,".egENSEMBL2EG"))

IDs=as.list(my.ENS2EG)

#Create two lists, one for the DEGs and one for the GOs
dewg = list()
gowg = list()

# A loop that goes through the DEG files we created earlier
for(i in 1:length(yy)){ #as many clusters as we have
  dewg[[i]] = xy[[i]] #Read the table
  ex = sapply(dewg[[i]], function(x) exists(x, my.ENS2EG)) #Which genes have an ENTREZ?
  dewg[[i]] = dewg[[i]][ex] #Only those genes
  dewg[[i]] = unlist(IDs[dewg[[i]]]) #The ENTREZ IDs
  gowg[[i]] = goana(dewg[[i]], species = my.sp, universe = unlist(IDs[rownames(Wdata)])) #The GO analysis
}

goplots=list()
goterms=list()

for (i in 1:length(yy)) {
  toplot=topGO(gowg[[i]], n= 50, ontology = c("BP"))
  
  #Change to character and then back to factor, to keep the order from TopGO
  toplot$Term = as.character(toplot$Term)
  toplot$Term = factor(toplot$Term, levels = unique(toplot$Term))
  
  goterms[[i]]=toplot
  
}

detach("package:limma", unload=TRUE)
# We can save the GO data
# save(assign(paste0(project_name, "_WGCNA_GO"), gowg), file = paste0("./data/",project_name,"_WGCNA_GO.rda"))
# save(assign(paste0(project_name, "_WGCNA_GOterms"), gowg), file = paste0("./data/",project_name,"_WGCNA_GOterms.rda"))

#TO make the acutal plot we're having, we combine all the GOs
g.plot = data.frame()
topterms = data.frame()

for (i in 1:length(goterms)) {
  topterms = goterms[[i]][1:5,]
  topterms$cluster = i
  g.plot = rbind(g.plot, topterms)
}

g.plot$cluster = as.factor(g.plot$cluster)

my.terms = nchar(as.character(g.plot$Term))>45
my.gterms = strtrim(g.plot$Term, 45)
my.gterms[my.terms] = paste0(my.gterms[my.terms],"...")

ggplot(g.plot, aes(x=Term, y=-log10(P.DE), fill=cluster)) +
    geom_point(aes(shape = cluster), size = 3) +
    theme(axis.text.x = element_text(angle = 90, hjust = 1)) +
    labs(x="GO Term", y="-log10 p. value") +
    scale_x_discrete(labels=my.gterms) +
    scale_fill_manual(name = "Module",
                      labels = yy,
                      values = yy) + 
    scale_shape_manual(name = "Module",
                       labels = yy,
                       values = rep(c(21,22,23,24,25),20)) + coord_flip()
```

And here the report of each of the modules

```
# Go trhough the modules
for(i in 1:length(xx)){
  
  # A table is done, where we put the genes names that are making-up the module, in rows of 10. We fill the las row with empty spaces
  ktable1 = data.frame( matrix( c(gnames[xy[[i]],2],rep(" ", 10-length(gnames[xy[[i]],2])%%10 )), ncol = 10, byrow = T ) )
  colnames(ktable1) = rep(".", 10)
  
  # Another table, where we put the top 10 GO terms, in two columns
  ktable2 = data.frame( matrix( as.character(goterms[[i]][1:10,1]), ncol = 2, byrow = F ) )
  colnames(ktable2) = rep(".", 2)
  
  # We make a plot in which we show the mean expression level of the co-expression module
  
  toplot = data.frame(p.Wdata@reductions[[my.dr]]@cell.embeddings)
  
  myplot = ggplot(toplot[order(p.MEList$averageExpr[,i]),],
                   aes_string(x=colnames(toplot)[1], y=colnames(toplot)[2])) +
      geom_point(aes_string(color=p.MEList$averageExpr[order(p.MEList$averageExpr[,i]),i])) +
      scale_size(range = c(1, 1)) +
      theme_void() +
      scale_colour_gradientn(colours = c("gray90", "gray90", yy[i], yy[i])) +
      labs(colour=levels(as.factor(dynamicColors))[i])
  
  mynet=mynets[[i]]
  
  mynetplot = ggnet2(mynet,
                         mode = "fruchtermanreingold",
                         layout.par = list(repulse.rad=network.size(mynet)^1.1,
                                           area=network.size(mynet)^2.3), # Give space in the middle
                         node.size = get.vertex.attribute(mynet,'membership'), max_size = 22, #The size of the nodes
                         node.color = levels(as.factor(dynamicColors))[i], # The color of the module
                         edge.size = "weight",
                         edge.color = "black",
                         edge.alpha = get.edge.attribute(mynet,'weight01'),
                         ) +
    theme(legend.position="none") +
    geom_label(aes(label=gnames[network.vertex.names(mynet),2]),
               fill = levels(as.factor(dynamicColors))[i],
               alpha = 0.5,
               color=lcols[i],
               fontface = "bold")
  
  # We put our mean expression plot toghether with the network plot
  grid.arrange(grobs=list(myplot,mynetplot), nrow=2)
  plot.new()
  dev.off()
  cat("\n")
  # A table showing the number, color and size of the module
  print(knitr::kable(data.frame(module=i, color=names(table(dynamicColors))[i], size=unname(table(dynamicColors)[i]))))
  cat("\n")
  # The other two tables we just made
  print(knitr::kable(ktable1))
  cat("\n")
  print(knitr::kable(ktable2))
  cat("\n")

}
```

NA

| module | color | size |
| --- | --- | --- |
| 1 | black | 105 |

NA

| . | . | . | . | . | . | . | . | . | . |
| --- | --- | --- | --- | --- | --- | --- | --- | --- | --- |
| Cstdc5 | Krt15 | Krtdap | Krt14 | Lgals7 | Dmkn | Sfn | Perp | Fgfbp1 | Krt5 |
| Krt17 | Mt2 | Mgst1 | Epcam | Them5 | Gjb2 | Wnt6 | Bcam | Aqp3 | Lmo1 |
| Dsp | Serpinb10 | Tfap2b | Kremen2 | Endou | Gata3 | Cldn6 | Meis2 | Irx2 | Col17a1 |
| Nrarp | Trim29 | Barx2 | Wnt4 | Gjb6 | Trp63 | Tubb3 | Pkp3 | Hspb8 | Ch25h |
| Cdh1 | Pdlim1 | Pltp | Wnt7b | Urah | Chchd10 | Bicdl2 | Tfap2a | Gchfr | Wnt10a |
| Plet1 | Anxa9 | Pkp1 | Car12 | Ly6g6d | Mapk13 | Spns2 | Ces2g | Dpp4 | Irx4 |
| Jag1 | Ppp1r14c | Egfr | Klc3 | Serpinb5 | Ptprf | Anxa8 | Rdh12 | Rab38 | Gm13219 |
| Wnt3 | St14 | Lypd6b | Atp8b1 | Adh1 | Prss27 | Ifi202b | Zfp296 | Esrp2 | Irf6 |
| Lamb3 | Cgn | Ddr1 | Fras1 | Fzd10 | Esrp1 | Tfap2c | Pard6b | Trim2 | Etl4 |
| Cebpa | Akap17b | Fzd6 | Ckmt1 | Nipal1 | Hoxb4 | Igsf9 | Hotairm1 | Prxl2a | Gprc5a |
| Frem2 | Tmem184a | Fermt1 | Lamc2 | Itga3 |  |  |  |  |  |

NA

| . | . |
| --- | --- |
| epidermis development | epidermal cell differentiation |
| morphogenesis of an epithelium | canonical Wnt signaling pathway |
| tissue morphogenesis | intermediate filament organization |
| skin development | multicellular organismal water homeostasis |
| epithelium development | cell-cell signaling by wnt |

NA

NA

| module | color | size |
| --- | --- | --- |
| 2 | blue | 226 |

NA

| . | . | . | . | . | . | . | . | . | . |
| --- | --- | --- | --- | --- | --- | --- | --- | --- | --- |
| Cxcl14 | Crabp1 | Chodl | Postn | Fn1 | Stmn2 | Tnmd | Egfl6 | Tpm2 | Tac1 |
| Mmp11 | Igfbp2 | Aspn | Wif1 | Ctsk | Prrx1 | Scx | Tmeff2 | Map1b | Vcan |
| Calb1 | Rbp1 | Vcam1 | Rdh10 | Tbx18 | Ngfr | Kcnk2 | Pdlim3 | Sbspon | Col8a1 |
| Col6a3 | Thy1 | Ebf2 | Prrx2 | Sdc2 | Lmna | Cald1 | Dpt | a | Tmem30b |
| Cpxm2 | Twist1 | Rspo1 | Hmgn3 | Tril | Hand2os1 | Irx1 | En1 | Serpinc1 | Col7a1 |
| Tmem119 | Epha3 | Zeb2 | Nkd2 | Col23a1 | Irx3 | Tmem132c | Pax1 | Col14a1 | Gstt1 |
| Efna5 | H2bu2 | Svbp | Rasl11b | Cdo1 | Apela | Gsta4 | Gm14964 | Plxna4 | C1qtnf2 |
| Adam33 | Pcdh11x | Dkk1 | Plpp3 | Bex1 | Dpysl3 | Bex4 | Cntfr | Ltbp1 | Creb3l1 |
| Vegfd | Hand2 | Stra6 | Uchl1 | Rnd3 | Nt5e | Fez1 | Eva1b | Fam174b | Lix1 |
| Inhba | Ppp1r14a | Ptchd4 | Dnm3 | Adamts5 | Mkx | Pdgfc | Hic1 | Olfml3 | Thsd7a |
| Itih5 | Amot | NA | Thbs2 | Sema3a | Col5a1 | Sp5 | Vstm2b | Septin7 | Pdpn |
| Fap | Lsamp | Frem1 | Robo1 | Rhoj | Cttnbp2 | Rgs17 | Ptk7 | Tia1 | Spats2l |
| Ccn4 | Grem2 | C130021I20Rik | Cdh2 | Alx3 | Pdlim4 | Septin11 | Pianp | Fjx1 | Adgrg2 |
| C1qtnf7 | Rgs7bp | Pdgfrb | Bmp3 | Ism1 | Tpbg | Rarres2 | Mex3b | Insyn1 | Scara3 |
| Igsf10 | Fnbp1l | Pnck | Prr16 | Fgf10 | Smarca2 | Emx2 | Ptprd | Chst2 | Cbr3 |
| Tmem26 | Mme | Fgf18 | Lama2 | Palm | Fat4 | Nnmt | Tex15 | Cacng4 | Iglon5 |
| Rai14 | Gpc4 | Homer2 | Bend5 | Fndc1 | Zfp536 | Axin2 | Mylip | Ednra | Sema5a |
| Rimklb | Zfp503 | Lrrn3 | Fblim1 | Frmd4a | Cpne5 | Lmx1b | Cnih2 | Nradd | Serping1 |
| Sbk1 | Cd248 | Gm10371 | Klhdc8b | Cdc42ep5 | Wbp1 | NA | Kcns3 | Kcna4 | Cox20 |
| Plppr3 | Tgfbr3 | Ephb2 | Cacna2d3 | Itga8 | Ccdc136 | Gulp1 | St8sia2 | Slit3 | Tspan11 |
| Stox2 | Scube2 | Kirrel3 | Lin7a | Wnt11 | Enox1 | Dtx4 | Cplx2 | Lrig1 | Frmd6 |
| Egflam | Shd | Adamts2 | Unc5d | Cyth3 | Slc7a10 | Adam22 | Fbn1 | Samd14 | Dchs1 |
| Pappa2 | Grik5 | Vstm4 | Hunk | NA | Cspg5 |  |  |  |  |

NA

| . | . |
| --- | --- |
| anatomical structure morphogenesis | system development |
| multicellular organism development | animal organ morphogenesis |
| regulation of neuron projection development | developmental growth involved in morphogenesis |
| circulatory system development | positive regulation of neuron projection development |
| multicellular organismal process | regulation of plasma membrane bounded cell projection organization |

NA

NA

| module | color | size |
| --- | --- | --- |
| 3 | brown | 223 |

NA

| . | . | . | . | . | . | . | . | . | . |
| --- | --- | --- | --- | --- | --- | --- | --- | --- | --- |
| S100a8 | S100a9 | Cd74 | H2-Aa | Cstdc4 | H2-Ab1 | Ccl4 | H2-Eb1 | Pf4 | Ccl3 |
| Klrd1 | Plac8 | C1qb | Prtn3 | Srgn | Cxcl2 | Cd52 | C1qa | C1qc | S100a4 |
| Coro1a | Selenop | Hp | Rac2 | Rgs1 | Mgl2 | Fcrls | Cx3cr1 | Wfdc21 | Wfdc17 |
| Il1b | H2-D1 | Mrc1 | F13a1 | Ifitm6 | Laptm5 | Cd53 | Napsa | Rgs2 | Pglyrp1 |
| Bcl2a1b | Emb | Il1r2 | H2-K1 | Fxyd5 | Ccl2 | Ccl6 | Ms4a7 | Cd83 | Psmb8 |
| Cd68 | Mcemp1 | Capg | Cfp | Plbd1 | Clec10a | H2-DMa | Otulinl | Lcp1 | Trf |
| Evi2a | Tnni2 | Ptpn18 | F10 | H2-DMb1 | Gmfg | Lat2 | Il13ra1 | Fcgr2b | C3ar1 |
| Cxcl3 | Xlr | Tlr2 | Ighm | NA | Cytip | Bst2 | Rtp4 | Osgin1 | C5ar1 |
| Ncf4 | Rgs18 | Hcls1 | Gm2a | Glipr1 | Ramp1 | Rgs10 | Fermt3 | Cd79b | H2-Oa |
| Gpsm3 | Fgl2 | Pmaip1 | Dusp5 | Hck | Clec4d | AI662270 | Cd14 | Cd44 | Tpd52 |
| Apbb1ip | Fmnl1 | Hpgds | Lgals9 | Vsir | Dok2 | Lpxn | Selplg | Tuba4a | Mndal |
| Ptprc | Tm6sf1 | Nrros | Arhgap15 | Fyb | Sp110 | Gch1 | Hmox1 | Apobec3 | Dock2 |
| Bin2 | Tmem106a | Milr1 | Itgb2 | B4galnt1 | Ehd4 | Bcl2a1d | Osm | Sp140 | Pirb |
| Il18rap | Cyp4f18 | Myo1g | Oasl2 | Pak1 | Lyn | Sh2d1b2 | Clec2i | Ifi211 | Ptpro |
| Tnfaip8l2 | Mcub | Fam107b | Hfe | Gas7 | Gpr65 | Ltb4r1 | Zc3hav1 | Hsd11b1 | Ang |
| Lcp2 | Stom | Plaur | Slc11a1 | Pkib | Lpcat2 | Slamf9 | Casp1 | P2ry12 | Kcnn4 |
| Ifi203 | Ly9 | Smim3 | Il4ra | Bcl2a1a | Irf8 | Bank1 | Cd300lf | Emilin2 | Arhgap45 |
| Csf3r | Atp2a3 | Slc9a3r1 | 5430427O19Rik | Syngr2 | Adgre5 | Cysltr1 | Clec5a | Trpv2 | Gpr137b-ps |
| Pira2 | Ahr | Klhl6 | Tifa | Siglece | Gm14548 | Adssl1 | Arhgap9 | Phf11b | Ppfia4 |
| H2-T23 | Fam129a | Aph1c | Fgd2 | Rab32 | Tifab | Nckap1l | Prkcb | Blnk | Ccrl2 |
| Bmp2k | Gbp7 | Adcy7 | Vav1 | Ptpn7 | Rab7b | Tmem37 | Ptprj | Skap2 | Sirpa |
| Hhex | Lrmp | Fes | Ccr5 | Atp1a3 | Cd33 | Rassf4 | Mir22hg | Slc15a3 | Adap2os |
| Hacd4 | Psmb9 | Nlrp3 |  |  |  |  |  |  |  |

NA

| . | . |
| --- | --- |
| immune response | positive regulation of immune response |
| immune system process | positive regulation of immune system process |
| innate immune response | regulation of immune system process |
| defense response | immune effector process |
| regulation of immune response | leukocyte activation |

NA

NA

| module | color | size |
| --- | --- | --- |
| 4 | cyan | 58 |

NA

| . | . | . | . | . | . | . | . | . | . |
| --- | --- | --- | --- | --- | --- | --- | --- | --- | --- |
| Lyz2 | Ifi27l2a | Fcer1g | Tyrobp | Ctss | Ccl9 | Alox5ap | Ctsc | Ms4a6c | Lst1 |
| Clec4a2 | Ccr2 | Fcgr3 | Ms4a6b | Aif1 | Ly86 | Csf1r | Clec4n | Ms4a6d | Clec4a1 |
| Spi1 | Pld4 | Adgre1 | Trem2 | Slfn2 | Clec12a | Gm49339 | Fcgr1 | Ccr1 | Plek |
| Ms4a4c | Cd300c2 | Clec4a3 | AB124611 | Cd48 | Unc93b1 | Cybb | P2ry6 | Apoc2 | Rnase6 |
| Clec7a | Ifi207 | Igsf6 | Ifi204 | Ptpn6 | Mpeg1 | Cd86 | Ptafr | Epsti1 | Apobec1 |
| Tnfrsf13b | Cyth4 | Ncf2 | Lrrc25 | Myo1f | Ebi3 | Irf5 | Pilra |  |  |

NA

| . | . |
| --- | --- |
| defense response | antigen processing and presentation of peptide antigen |
| immune response | myeloid leukocyte activation |
| immune system process | response to other organism |
| innate immune response | response to external biotic stimulus |
| adaptive immune response | response to biotic stimulus |

NA

NA

| module | color | size |
| --- | --- | --- |
| 5 | green | 151 |

NA

| . | . | . | . | . | . | . | . | . | . |
| --- | --- | --- | --- | --- | --- | --- | --- | --- | --- |
| Hist1h2ap | Ptma | Tubb5 | H2az1 | Stmn1 | Hmgb2 | Tuba1b | Ube2c | Xist | Cenpa |
| Pclaf | Birc5 | Arl6ip1 | Cdc20 | Rrm2 | H1f0 | Cks2 | Top2a | H2ax | Ube2s |
| Ccnb1 | Tubb4b | Hmgn2 | Cenpf | H1f10 | Calm2 | Jpt1 | Lockd | Nusap1 | H2az2 |
| Cdca3 | Cdca8 | Ccnb2 | Lsm6 | Col4a2 | Slc24a5 | Smc2 | Cenpe | Spc25 | Smc4 |
| Pbk | Tpx2 | Cdk1 | Ccna2 | Tubb6 | Dek | Tyms | Tk1 | Cbx3 | Hmgb3 |
| Tacc3 | Spc24 | Kif23 | Hmmr | Ccdc34 | Lig1 | Tafa4 | Tuba1c | Ptms | Nucks1 |
| Slbp | Dut | Bub3 | Tmpo | Racgap1 | Mki67 | Prc1 | Knstrn | Lmnb1 | Hoxc10 |
| Cit | Ccne1 | Aurkb | Spock3 | Incenp | Kif20b | Depdc1a | Psrc1 | Sms | Cdkn3 |
| Fxyd6 | Pimreg | Tcf19 | Plk1 | Kif22 | Slc29a1 | Gm47283 | Asf1b | Rpgrip1 | Pcna |
| Fhl1 | Cdkn2c | Mcm5 | Kif20a | Hoxa9 | Nsg1 | Aurka | Plk4 | H2ac8 | Mcm2 |
| Aspm | Mcm3 | Dlgap5 | Nuf2 | Knl1 | Syne2 | 1500009L16Rik | Neil3 | Mis18bp1 | Melk |
| Esco2 | Syce2 | Gmnn | Ncapd2 | Mid1 | Nde1 | Hells | Kifc1 | Shcbp1 | Supt16 |
| Kif2c | Sgo2a | Chaf1b | Ckap2l | Mxd3 | H4c9 | Fzd4 | Sapcd2 | Pik3r3 | Ndc80 |
| Uhrf1 | Gins2 | Ckap2 | Anln | Cdc25c | H1f4 | Bub1b | Gli1 | Kpna2 | Ube2t |
| Hs6st2 | Rad51 | Pkmyt1 | Lnpk | Ncapg | Clasp2 | Cep89 | Bub1 | Gas2l3 | Wdhd1 |
| Atad2 |  |  |  |  |  |  |  |  |  |

NA

| . | . |
| --- | --- |
| cell cycle | chromosome organization |
| cell cycle process | chromosome segregation |
| cell division | nuclear chromosome segregation |
| mitotic cell cycle | sister chromatid segregation |
| mitotic cell cycle process | nuclear division |

NA

NA

| module | color | size |
| --- | --- | --- |
| 6 | greenyellow | 70 |

NA

| . | . | . | . | . | . | . | . | . | . |
| --- | --- | --- | --- | --- | --- | --- | --- | --- | --- |
| Anxa1 | Il22 | Mzb1 | Prr29 | Prr11 | Crip1 | Actg1 | Cdkn1a | Nrgn | Car2 |
| Arg1 | Cd4 | Nfkbia | Ptgs2 | Gpx1 | Akr1c13 | Serpina3g | Dusp1 | Zap70 | Tspan13 |
| Zfp36 | Cited2 | Tspo | Ctsd | Rrad | Cnn2 | Gsn | Lat | Sdf2l1 | Tox2 |
| Ifi30 | Tgfb1 | Cited4 | Msn | Slc7a11 | Macroh2a1 | Ikzf2 | Actr3 | Septin1 | Cxcl16 |
| Ier5 | Serp1 | Ahnak | Hexb | Cmtm7 | Asb17 | Sec11c | Ifrd1 | Rap1b | Cdk2ap2 |
| Pik3r1 | Ifngr1 | Gsto1 | Tec | Dnase2a | Lrrfip1 | Arl4c | Zyx | Samd10 | Il17re |
| Ly6a | Hpcal1 | Samhd1 | Trim8 | Espn | Cyfip2 | Jmjd1c | Myb | Fam124b | Kdm7a |

NA

| . | . |
| --- | --- |
| immune system process | regulation of calcium ion transport |
| response to cytokine | regulation of immune system process |
| establishment of protein localization to organelle | regulation of cellular process |
| response to stress | response to interferon-gamma |
| tissue regeneration | regulation of cellular localization |

NA

NA

| module | color | size |
| --- | --- | --- |
| 7 | grey60 | 40 |

NA

| . | . | . | . | . | . | . | . | . | . |
| --- | --- | --- | --- | --- | --- | --- | --- | --- | --- |
| Tmsb4x | Col1a1 | Tmsb10 | Col3a1 | Col1a2 | Sfrp2 | Vim | Mdk | Igfbp4 | Ubb |
| Gas1 | Mfap4 | Bgn | Gas2 | Mfap2 | Col6a2 | Fbln5 | Marcksl1 | Sox11 | Fstl1 |
| Cxcl12 | Tcf4 | Nfib | Fbn2 | Il11ra1 | Runx1t1 | Cdh11 | Zfp36l1 | Pdgfra | Fgfr1 |
| 6330403K07Rik | Loxl1 | Wnt5a | Tnfaip6 | Mmp14 | Pfn2 | Hoxd12 | Macroh2a2 | Gm9844 | Clmp |

NA

| . | . |
| --- | --- |
| head development | digestive tract development |
| skeletal system development | appendage morphogenesis |
| central nervous system development | limb morphogenesis |
| brain development | regulation of cell migration |
| central nervous system neuron development | anatomical structure morphogenesis |

NA

NA

| module | color | size |
| --- | --- | --- |
| 8 | lightcyan | 50 |

NA

| . | . | . | . | . | . | . | . | . | . |
| --- | --- | --- | --- | --- | --- | --- | --- | --- | --- |
| Cma1 | Mcpt8 | Cpa3 | Tpsb2 | Hdc | Prss34 | Tpsg1 | Ftl1 | Cd200r3 | Cebpb |
| B2m | Cyp11a1 | Ly6e | Fth1 | Gata2 | Il1rl1 | Smpx | Sat1 | Il4 | Hs3st1 |
| Psap | Cyba | Pim1 | Il13 | Rab27b | Metrnl | Tph1 | Tesc | Cd200r4 | Cst7 |
| Alox5 | Slc45a3 | Adora3 | Snx2 | Gadd45a | Prkcq | Nr4a2 | Plin2 | Skap1 | Gyg |
| Tiparp | Cela1 | Tpsab1 | Gstt2 | Klk8 | Kit | Slc7a5 | Man2b1 | Gnaz | Ubash3b |

NA

| . | . |
| --- | --- |
| inflammatory response | lysosome localization |
| positive regulation of mast cell activation involved in immune response | mast cell degranulation |
| positive regulation of mast cell degranulation | mast cell activation involved in immune response |
| regulation of proton transport | mast cell mediated immunity |
| positive regulation of mast cell activation | positive regulation of leukocyte degranulation |

NA

NA

| module | color | size |
| --- | --- | --- |
| 9 | lightgreen | 31 |

NA

| . | . | . | . | . | . | . | . | . | . |
| --- | --- | --- | --- | --- | --- | --- | --- | --- | --- |
| S100a11 | Anxa2 | Gja1 | Dlx2 | Bcl11b | Fst | Ass1 | Calm4 | Sbsn | Tnfrsf19 |
| Cdh3 | Acaa2 | Fzd10os | 1110028F18Rik | Sox15 | Wnt16 | Megf6 | Limk2 | Ank3 | Svil |
| Cldn1 | Mpped1 | Ociad2 | Ltbp4 | Sult5a1 | Susd4 | Ccdc3 | Efcab15 | Tom1l1 | Dlx4os |
| Fam83f |  |  |  |  |  |  |  |  |  |

NA

| . | . |
| --- | --- |
| keratinocyte proliferation | subpallium development |
| forebrain neuron fate commitment | odontogenesis of dentin-containing tooth |
| skin development | regulation of hair follicle development |
| cell communication by electrical coupling | hair cycle process |
| epidermis development | hair follicle development |

NA

NA

| module | color | size |
| --- | --- | --- |
| 10 | lightyellow | 22 |

NA

| . | . | . | . | . | . | . | . | . | . |
| --- | --- | --- | --- | --- | --- | --- | --- | --- | --- |
| Dct | Pmel | Ptgds | Mlana | Ednrb | Plp1 | Trpm1 | Gpnmb | Npy | Foxd3 |
| Syngr1 | Gsta1 | Pax3 | Tyr | Slc45a2 | Mcoln3 | Bace2 | Slc26a7 | Sox10 | Insc |
| Phyhipl | Nkain4 |  |  |  |  |  |  |  |  |

NA

| . | . |
| --- | --- |
| pigmentation | secondary metabolite biosynthetic process |
| developmental pigmentation | pigment metabolic process |
| melanin biosynthetic process | secondary metabolic process |
| pigment biosynthetic process | phenol-containing compound biosynthetic process |
| melanin metabolic process | phenol-containing compound metabolic process |

NA

NA

| module | color | size |
| --- | --- | --- |
| 11 | magenta | 85 |

NA

| . | . | . | . | . | . | . | . | . | . |
| --- | --- | --- | --- | --- | --- | --- | --- | --- | --- |
| Krt4 | Krt19 | Asprv1 | Cnfn | Krt6a | Krt1 | Psca | Hspb1 | Upk1b | Sprr1a |
| Ly6d | Mal | S100a14 | Krt7 | Cldn3 | Tacstd2 | Cldn4 | Cldn23 | Krt8 | Slc2a3 |
| Krt18 | Dsc2 | Upk3bl | Gm16136 | Rab25 | Ppl | Lypd3 | Aldh3b2 | Lor | Mab21l4 |
| Capns2 | Gng13 | Pdzk1ip1 | Areg | Sult2b1 | Grhl1 | Clic3 | Ovol1 | Fam25c | Map3k6 |
| Mall | Paqr6 | Snx31 | Ly6g6e | Nectin4 | Dsc3 | Paqr5 | Jup | Gpr87 | Grhl3 |
| Zfp750 | Sp6 | Scel | Rhov | Tmem40 | Spint1 | Tmem125 | Nebl | Ablim1 | Adtrp |
| Aldh1a7 | Crb3 | Prss8 | Phldb3 | Csta1 | Dsg1a | NA | Tmem79 | Cers3 | Smtnl2 |
| Klf5 | Evpl | Bspry | Sytl1 | Gipc2 | Arap2 | Cst6 | Kctd1 | Prom2 | Mpzl3 |
| Eps8l2 | Ephx3 | Tent5b | Il1rn | Sptlc3 |  |  |  |  |  |

NA

| . | . |
| --- | --- |
| skin development | establishment of skin barrier |
| epidermis development | epithelial cell differentiation |
| keratinization | regulation of water loss via skin |
| epidermal cell differentiation | multicellular organismal water homeostasis |
| keratinocyte differentiation | water homeostasis |

NA

NA

| module | color | size |
| --- | --- | --- |
| 12 | midnightblue | 54 |

NA

| . | . | . | . | . | . | . | . | . | . |
| --- | --- | --- | --- | --- | --- | --- | --- | --- | --- |
| Dcn | mt-Atp6 | Dkk4 | Fabp5 | mt-Co3 | mt-Co1 | mt-Nd1 | mt-Cytb | mt-Co2 | Arhgdib |
| Sostdc1 | mt-Nd2 | Edar | Npnt | mt-Nd5 | Msx2 | Ier3 | Hspa1a | Syt1 | Irx6 |
| Jag2 | Spry1 | Notch3 | Syt13 | Cyb5a | Gdpd1 | Cygb | Tnfrsf12a | Emp2 | Hoxc13 |
| Ptprz1 | Rhbdl3 | Lypd6 | Bmp7 | Ddit4 | Cldn7 | Zfp185 | Tmem132a | Astn2 | Kctd11 |
| 1700093K21Rik | Nr2e3 | Hoxc8 | Mfhas1 | Nav2 | Foxo6 | Unc5b | F2rl1 | Prr5l | Narf |
| Sema3f | Cachd1 | Pard6g | 5730522E02Rik |  |  |  |  |  |  |

NA

| . | . |
| --- | --- |
| cellular respiration | purine nucleoside monophosphate metabolic process |
| ATP metabolic process | purine ribonucleoside monophosphate metabolic process |
| ATP synthesis coupled electron transport | ribonucleoside monophosphate metabolic process |
| purine ribonucleoside triphosphate metabolic process | purine nucleoside triphosphate metabolic process |
| ribonucleoside triphosphate metabolic process | electron transport chain |

NA

NA

| module | color | size |
| --- | --- | --- |
| 13 | pink | 89 |

NA

| . | . | . | . | . | . | . | . | . | . |
| --- | --- | --- | --- | --- | --- | --- | --- | --- | --- |
| Ccl5 | Actb | Serpinb1a | Ifitm1 | Cd7 | Ltb | Trbc2 | Nkg7 | Ptprcap | Trbc1 |
| Pfn1 | Tnfrsf9 | Tcf7 | Junb | Cd3g | Itm2b | Tagln2 | Apoc1 | Slc6a13 | Cd69 |
| Ckb | Klrb1b | Cd37 | Icos | Slc25a5 | Tagap | Sh3bgrl3 | Rora | Samsn1 | Cd160 |
| Ctsw | Ccr7 | Hcst | Il18r1 | Arpc1b | Flt3 | Cxcl10 | Xlr4b | Itgal | Itgb7 |
| Pou2f2 | Klf6 | Fam83a | Ikzf1 | Rasal3 | Xlr4a | Clic1 | Sla | Icam1 | Asb2 |
| Cxcr6 | Tnfrsf25 | Il2rg | Npl | Cd28 | Akr1b3 | Arl6ip5 | S100a13 | Crem | Xlr4c |
| Cd47 | Lta | Il7r | Mitf | Gpr183 | Rhoh | Serinc3 | Slc16a10 | Ecm1 | Entpd1 |
| Gpr171 | Parvg | Rnasel | Itk | Rnf130 | Tnfaip3 | Ctnnd2 | Csrnp1 | Zfp263 | Fam111a |
| Tmem154 | Dleu2 | Rbm38 | Runx1 | Rab37 | Irak2 | Abca3 | Esyt1 | Rassf2 |  |

NA

| . | . |
| --- | --- |
| immune system process | cell activation |
| T cell activation | leukocyte differentiation |
| lymphocyte activation | immune effector process |
| leukocyte activation | T cell differentiation |
| alpha-beta T cell activation | regulation of immune system process |

NA

NA

| module | color | size |
| --- | --- | --- |
| 14 | purple | 72 |

NA

| . | . | . | . | . | . | . | . | . | . |
| --- | --- | --- | --- | --- | --- | --- | --- | --- | --- |
| Gdf5 | Serpine2 | H3f3b | Hes1 | Htra1 | Dkk3 | Msx1 | Ccnd2 | Col8a2 | Nell2 |
| Ptn | Pthlh | Nrtn | Hoxd13 | Fxyd7 | Hoxa11os | Ccnd1 | Sdc1 | Foxp2 | Col12a1 |
| Fbln2 | Tgfb2 | Sema3c | Ntng1 | Creb5 | Lmo4 | Nrxn1 | Serpinf1 | Mecomos | Hhip |
| Erg | Hmcn1 | Lhx2 | Epha4 | Nrn1 | Mpped2 | Car14 | Hoxa10 | Frk | Hoxd11 |
| Tob1 | Pgrmc1 | Ssbp2 | Syt11 | Mn1 | Srsf12 | Axl | Serinc2 | Tmtc2 | Tgfb3 |
| 2700069I18Rik | Fam171b | Prdm16 | Sv2a | Pcdh10 | Aopep | Thsd4 | Kcnd2 | Car11 | Adgrl3 |
| Pbx1 | Plxdc2 | Erc2 | Igsf3 | Slit2 | Kcnmb4 | Foxp4 | Mllt3 | Ank2 | Dlg5 |
| Tenm3 | Bmp1 |  |  |  |  |  |  |  |  |

NA

| . | . |
| --- | --- |
| regulation of RNA metabolic process | regulation of nucleic acid-templated transcription |
| reproductive structure development | regulation of transcription, DNA-templated |
| reproductive system development | regulation of RNA biosynthetic process |
| gland development | RNA metabolic process |
| regulation of gene expression | nucleic acid-templated transcription |

NA

NA

| module | color | size |
| --- | --- | --- |
| 15 | red | 119 |

NA

| . | . | . | . | . | . | . | . | . | . |
| --- | --- | --- | --- | --- | --- | --- | --- | --- | --- |
| Acta2 | Actc1 | Myog | Tnnt2 | Tnni1 | Mylpf | Myl1 | Tnnt1 | Mymx | Igfbp3 |
| Vgll2 | Cbln2 | Car3 | Cryab | Rplp1 | Chrna1 | Mymk | Msc | Kcne1l | Tnnc2 |
| Ttn | Des | Ptgis | Cdh15 | Myod1 | Pgam2 | Atp2a1 | Myl4 | Tnnc1 | Il17b |
| Cck | Necab1 | Sln | Pdgfa | Gatm | Shisa2 | Six1 | Olfml2b | Celf2 | Tubb2b |
| Klhl41 | Mstn | Nes | Tppp3 | Septin4 | Jsrp1 | Hspb2 | Tmem200a | Spon1 | Pitx2 |
| Hspd1 | Tex14 | Rbm24 | Gpr37 | Rgs16 | Casq2 | Sypl2 | Fmo1 | Ccdc155 | Myf5 |
| Srpk3 | Nexn | Trim63 | Cited1 | Parm1 | Fitm1 | Neb | Srl | Myh3 | Eya2 |
| Mycl | Tmem35a | Lrrn1 | Unc45b | Rai2 | Dbx1 | Arpp21 | Ank1 | Tnik | Myl6b |
| Iffo1 | Fndc5 | Acta1 | Tcea3 | AW551984 | Mrln | Lurap1l | Camk2n1 | Tceal6 | Pnmal2 |
| Chrng | Dcx | Flnc | Pdlim7 | Ssc5d | Cap2 | Prkaca | Cox6a2 | Fam110b | Rtn2 |
| Epor | Ralgps2 | Shisal2a | Dll1 | Hdac11 | Casz1 | Traf3ip3 | Frmd4b | Pgm5 | Arhgdig |
| Dbndd1 | Itga7 | Bcl7a | Rhobtb1 | Mad2l2 | Tln2 | Rxrg | Vash2 | Megf10 |  |

NA

| . | . |
| --- | --- |
| muscle structure development | striated muscle contraction |
| muscle system process | skeletal muscle organ development |
| muscle organ development | muscle tissue development |
| striated muscle tissue development | skeletal muscle tissue development |
| muscle contraction | muscle cell development |

NA

NA

| module | color | size |
| --- | --- | --- |
| 16 | salmon | 65 |

NA

| . | . | . | . | . | . | . | . | . | . |
| --- | --- | --- | --- | --- | --- | --- | --- | --- | --- |
| Lgals1 | Dkk2 | Sox4 | Igfbp5 | Col6a1 | Tuba1a | Nnat | Nbl1 | Tpm1 | Foxp1 |
| Ogn | Marcks | Col4a1 | Crabp2 | Agtr2 | Osr2 | Bmp4 | Ly6h | Sulf1 | Fbln1 |
| Lox | Osr1 | Crlf1 | Nfia | Col5a2 | Csrp2 | Lhfp | Dclk1 | Basp1 | Epha7 |
| Ddah2 | Col26a1 | Mecom | Tle5 | Chd3 | Lpar1 | Csrp1 | Dbn1 | Bcl11a | Cacna1g |
| Angptl4 | Glipr2 | Ntf3 | Rbms3 | Adamtsl1 | Zfhx4 | S1pr3 | Cmtm3 | Robo2 | Cped1 |
| Ctxn1 | Tead2 | Aplp1 | Myh10 | Epb41l3 | Sncaip | Lhfpl2 | Podxl2 | Phldb2 | Pcdh18 |
| Myo1b | Nfatc4 | Mir100hg | Cacnb3 | Efnb3 |  |  |  |  |  |

NA

| . | . |
| --- | --- |
| renal system development | ureter development |
| urogenital system development | kidney development |
| cellular response to acid chemical | actomyosin structure organization |
| head development | anatomical structure morphogenesis |
| brain development | central nervous system development |

NA

NA

| module | color | size |
| --- | --- | --- |
| 17 | tan | 66 |

NA

| . | . | . | . | . | . | . | . | . | . |
| --- | --- | --- | --- | --- | --- | --- | --- | --- | --- |
| Mmp13 | Ibsp | Mmp9 | Rbp4 | Gpx3 | Dmp1 | Steap4 | Fam20c | 2200002D01Rik | Ddit4l |
| Maf | Fabp3 | Ffar4 | Frzb | Cp | AW112010 | Lpl | Clu | Foxq1 | Slpi |
| Irx5 | Slc36a2 | Serpini1 | Hand1 | Rasl11a | Cemip | Ocm | Enpp6 | Itgb3 | Hpgd |
| Entpd3 | Chst1 | B3galt2 | Gm19705 | Loxl4 | Bmp8a | Tcim | Agt | Lratd1 | Lgr6 |
| Ptges | Dlx5 | Dapk2 | Hmgcs2 | Podnl1 | Ank | 9130024F11Rik | Cdkn2b | Tnfaip2 | Ackr3 |
| Csf1 | Cds1 | Mgst2 | Gm10638 | Phex | Syna | Jam2 | Nos1ap | Adamts9 | Pla2g4a |
| Cd55 | Sp7 | Rcan2 | Plaat3 | A230083N12Rik | NA |  |  |  |  |

NA

| . | . |
| --- | --- |
| cellular lipid metabolic process | small molecule metabolic process |
| lipid biosynthetic process | copper ion transport |
| positive regulation of protein kinase C activity | foam cell differentiation |
| regulation of protein kinase C activity | macrophage derived foam cell differentiation |
| lipid metabolic process | regulation of macrophage derived foam cell differentiation |

NA

NA

| module | color | size |
| --- | --- | --- |
| 18 | turquoise | 308 |

NA

| . | . | . | . | . | . | . | . | . | . |
| --- | --- | --- | --- | --- | --- | --- | --- | --- | --- |
| Col2a1 | Meg3 | Col9a3 | Cdkn1c | Matn1 | Itm2a | Cytl1 | H19 | Dlk1 | Snorc |
| Lum | Col11a1 | Epyc | Col9a1 | Fos | Cthrc1 | Uts2b | Mgp | Col9a2 | Cnmd |
| C1qtnf3 | Serpinh1 | Jun | Cpe | Klf2 | Matn4 | Mest | Ifitm5 | Hapln1 | Mia |
| Gng11 | Comp | Col11a2 | Jund | Matn3 | Acan | Penk | Ihh | Ostn | Smpd3 |
| Cd24a | Sparc | Ndufa4l2 | Ucma | Higd1a | Egr1 | Ier2 | Scrg1 | Sgms2 | Lgals3 |
| Ebf1 | Cd9 | Fabp7 | Tsc22d1 | Tgfbi | Fdps | Pth1r | Col27a1 | Panx3 | Rcn3 |
| Ccn1 | Ecrg4 | Mif | Id3 | Fibin | Ppa1 | Hspa5 | Fmod | Pcp4 | H1f2 |
| Pcolce2 | Pcolce | S100b | Gpc3 | Dbi | Sox9 | Smoc2 | Msmo1 | Nfatc2 | Id1 |
| Neat1 | Lbhd2 | Klf4 | Enpp2 | Timp3 | Kdelr2 | Thbs1 | Gdf10 | Hsp90b1 | Serinc5 |
| Chad | Susd5 | Id2 | Ntm | Angptl1 | Ier3ip1 | Cd82 | Maged2 | Capn6 | Ctsz |
| Rian | Plod2 | Klk10 | Prelp | P4ha2 | Barx1 | Phlda1 | Slc16a3 | Rcn2 | Peg3 |
| 3830403N18Rik | Plagl1 | Gale | Chadl | Papss2 | Fkbp9 | Scd2 | Bambi | Col15a1 | Arsi |
| Anxa5 | Phlda2 | Bnip3 | Saa1 | Kcnq1ot1 | Fosb | Moxd1 | Kazald1 | Wwp2 | Pmp22 |
| Zbtb20 | Rspo3 | Bmp2 | Mvd | Hmgcs1 | Fkbp11 | Mgarp | Nog | Fam162a | Sdc4 |
| Sox5 | Cox4i2 | Serpina3n | Clec11a | Ugdh | Acat2 | Fgfr3 | Tspan4 | Mpz | Fam180a |
| Pam | Asb4 | Rgcc | Shox2 | Vkorc1 | Htra3 | Emilin1 | Tmem158 | Rtl3 | Pcsk6 |
| P4ha1 | Cdc42ep3 | Hmga2 | Mfge8 | Rflna | Bex2 | Tubb2a | Bcl2 | Timp1 | Kdelr3 |
| Dab2 | Nrk | Selenom | Arl4a | Dhx58os | Ldhb | Fbln7 | Fkbp10 | Tcf7l2 | Insig1 |
| Islr | Klhl13 | Srm | Auts2 | Snhg18 | Saa2 | Trps1 | Ndrg2 | Fzd9 | P3h3 |
| Ppic | Prss35 | Pdcd4 | Rgs3 | Dap | Smim14 | 2810403D21Rik | Nid2 | Calml3 | Idi1 |
| Foxc2 | Cyp51 | Slc1a3 | Alkbh1 | Bhlhe41 | Btbd3 | Pxdc1 | Cdh10 | Trim47 | Loxl3 |
| Steap1 | Pitx1 | Tmem97 | Wfdc12 | Tspan8 | S100a1 | Lrig3 | Sorbs2 | Cox6b2 | Il17d |
| Bmper | Pantr1 | Smoc1 | Foxc1 | Tent5a | Cspg4 | Tspan18 | Cpxm1 | Runx3 | Clic4 |
| Fam89a | Ccn2 | Sqle | 1700049E15Rik | Trib1 | Klf9 | Ung | Ddah1 | Spsb4 | BC006965 |
| Boc | Scin | Dusp14 | Flrt2 | Lpar4 | Mxra8 | Scd1 | Meltf | Angptl2 | Trabd2b |
| Efna1 | Id4 | Dhcr7 | Efcab10 | Bmp5 | 4930523C07Rik | Hmgcr | Aebp1 | Runx2 | Tshz2 |
| Tbx15 | Cpq | Cldn10 | Sfrp1 | Raph1 | Gem | Smim5 | Psma8 | Slc16a4 | Gm26532 |
| Tspan32 | Slc26a2 | Fgfr2 | Tox | Stk32b | Chil1 | Lmcd1 | Srpx | Rlbp1 | Sema3d |
| Phyh | Cyp26b1 | Aldoc | Vwc2 | Kcnk1 | Batf3 | Cpm | Fam81a | Pxylp1 | Crispld1 |
| 2610528A11Rik | Aacs | Ttll3 | Tgm2 | Fbp1 | Enpp1 | Ltbp3 | Car9 | Cgref1 | Prkg2 |
| 1700086L19Rik | Ppp2r2b | Uba7 | Nim1k | Scn1b | Zfp24 | Ooep | Lss |  |  |

NA

| . | . |
| --- | --- |
| connective tissue development | extracellular matrix organization |
| cartilage development | sterol biosynthetic process |
| ossification | steroid biosynthetic process |
| chondrocyte differentiation | extracellular structure organization |
| skeletal system development | alcohol biosynthetic process |

NA

NA

| module | color | size |
| --- | --- | --- |
| 19 | yellow | 213 |

NA

| . | . | . | . | . | . | . | . | . | . |
| --- | --- | --- | --- | --- | --- | --- | --- | --- | --- |
| Ccl21a | Cldn5 | Ifitm3 | Gja4 | Fabp4 | Plvap | Emcn | Cdh5 | Lyve1 | S100a16 |
| Cav1 | Meox1 | Esam | Ctla2a | Cd34 | Aplnr | Mmrn1 | Gpihbp1 | Igf1 | Ecscr |
| Edn1 | Icam2 | Tm4sf1 | Cotl1 | Pecam1 | Gm16104 | Kcne3 | Sox17 | Col18a1 | Cavin2 |
| Flt1 | Vwf | Madcam1 | Nrp2 | Gimap6 | Myo1c | Rasip1 | Eng | Nts | Esm1 |
| Cd93 | Plxnd1 | Adm | Tmem255a | Crip2 | Kdr | Gap43 | Smagp | Myct1 | Lmo2 |
| Meox2 | Tie1 | Ly6c1 | Tpm4 | Stab1 | Gimap4 | Prcp | Fam167b | BC028528 | Klhl4 |
| Dusp2 | Depp1 | Gm525 | Fkbp1a | Dok4 | Cavin3 | Tinagl1 | Akap12 | Cd36 | Angpt2 |
| F11r | Clec14a | Tc2n | Tmem252 | Lpar6 | Trp53i11 | Mfng | Gngt2 | Flt4 | Grrp1 |
| Grap | Robo4 | Pcdh17 | Arhgap18 | Myzap | Mmrn2 | Prox1 | Apln | Mcam | Apold1 |
| Reln | Afap1l1 | Tmem88 | Clec1a | Rbpms | Lhx6 | Sox7 | Gm45837 | Adgrf5 | Hoxb9 |
| Nr2f2 | Cmtm8 | Gimap1 | Rasgrp2 | C130074G19Rik | Tek | Gm13889 | Gimap5 | Nectin2 | Pdgfb |
| Sult1a1 | Ptprb | Crmp1 | Hoxb6 | Ccl21d | Hlx | Pgf | Ushbp1 | Gpm6a | Ets1 |
| Arhgef28 | Bcl6b | Kank3 | Stc1 | Cav2 | Vamp5 | Dll4 | Tspan15 | AU021092 | Procr |
| Ipo11 | Palmd | Lamb1 | N4bp3 | F2r | Tfpi | 4930542C12Rik | Adgrl4 | Fhl2 | Bok |
| Hdac7 | Enpp3 | Sipa1 | Podxl | Npdc1 | Ephb1 | Ldb2 | Arap3 | Cdh13 | Thsd1 |
| Ece1 | Cd40 | Lck | Cpa1 | Acvrl1 | Ndrg1 | Npr1 | Marchf11 | Bik | Cd1d1 |
| Hid1 | Tnfaip8l1 | Prkch | Ppp1r16b | Cracr2b | Vwa1 | Arhgef15 | Fam43a | Itga6 | C1qtnf9 |
| Fgd5 | Ehd2 | Kitl | Adcy10 | Epas1 | Cda | Cnrip1 | Cavin1 | Lama4 | Arhgap31 |
| Dysf | Adam15 | Apoa2 | Cd38 | Prrg3 | Nedd9 | Mmp15 | Abi3 | Arhgap29 | Fam171a2 |
| Ccdc149 | Rftn1 | Hbegf | Slc16a13 | Pcdh1 | Filip1 | Cngb1 | Scarf1 | Kcna5 | Calcrl |
| Nova2 | Tmod2 | Arhgef7 | Tspan12 | Zfp979 | Ccm2l | Hoxd9 | Tcf15 | Dusp3 | Sema6d |
| Exoc3l4 | Snrk | Sh3tc1 |  |  |  |  |  |  |  |

NA

| . | . |
| --- | --- |
| vasculature development | circulatory system development |
| cardiovascular system development | endothelium development |
| angiogenesis | anatomical structure formation involved in morphogenesis |
| blood vessel development | tube morphogenesis |
| blood vessel morphogenesis | endothelial cell differentiation |

NA
