## Supplement 2 for "Assessing evolutionary and developmental transcriptome dynamics in homologous cell types"

WGCNA preservation, E15 as reference


Code 

- Show All Code
- Hide All Code
- Download Rmd

### WGCNA preservation, E15 as reference

###### Christian Feregrino

Date: 23.12.20

This script tries to test the preservation of network modules in different samples. It follows the tests presented in Langfelder et al. 2011 and the documentation and examples provided by the authors in different documents

#### Preservation of modules in other samples

Here, we use a network, and set of modules (all) calculated from one sample. We test the preservation of these modules in other samples and can compare the preservation statistics across modules.

We need several packages and load in the datasets.

Its important that the data sets share all their genes, so that we are able to make comparisons.


So we first take out test samples, and subset only for 1-to-1 orthologs


```
setwd("~/projects/scMmGg_Oct20/WGCNA/")

# placeholder for instructions from gigher level pipelines

my.121 = read.csv("./../data/Mm_121_Gg_ENS101.txt", fill = T, header = T, colClasses = c("character", "character", "character","integer"))
my.121=my.121[which(my.121$Chicken.homology.type == "ortholog_one2one" & my.121$Chicken.orthology.confidence..0.low..1.high. == 1 ),]

my.ortho = my.121[,c(1,2)]

# Datasets we're using

my.test = list()
my.test[[1]] = readRDS("./data/E9_subsample_pseudocells_171220.rds")
my.test[[2]] = readRDS("./data/E10_subsample_pseudocells_171220.rds")
my.test[[3]] = readRDS("./data/E11_pseudocells_291020.rds")
my.test[[4]] = readRDS("./data/E13_pseudocells_291020.rds")
my.test[[5]] = readRDS("./data/E18_pseudocells_171220.rds")
my.test[[6]] = readRDS("./data/HH21_pseudocells_291020.rds")
my.test[[7]] = readRDS("./data/HH24_pseudocells_291020.rds")
my.test[[8]] = readRDS("./data/HH25_pseudocells_171220.rds")
my.test[[9]] = readRDS("./data/HH27_pseudocells_291020.rds")
my.test[[10]] = readRDS("./data/HH29_pseudocells_171220.rds")
my.test[[11]] = readRDS("./data/HH31_pseudocells_181220.rds")

Wtest_names = c("E9","E10","E11", "E13", "E18","HH21","HH24","HH25","HH27","HH29","HH31")
my.ortho.sp = c(rep(1,5),rep(2,6))

for (i in 1:length(my.test)) {
  my.test[[i]] = subset(my.test[[i]], features = my.ortho[,my.ortho.sp[i]])
}


# What are the datasets to test WGCNA modules in?
Wtest = my.test

# Project name, will come out in files and folders
project_name="E15"

# The reference WGCNA data.
WGCNA_data = readRDS("data/E15_WGCNA_data.rds")
```


Now, we need to pre-process the data.

We need the test data sets in a list, as well as names to identify the samples. For the reference dataset, we need the WGCNA data in a list form. The list must contain at least:

- An expression matrix cells/genes like that used in the WGCNA module calculation
- The dynamic mods vector resulting from the module calculations

We subset the modules to have genes that are present in all samples. This is a drawback of the method. In order to make comparisons, we need variance in the data. If the gene is not expressed in one of the data sets, at all, there is no variance and the test can’t be made, but the rest of the module might still be present.

Once we have module genes that are expressed in all the data sets, we subset the data for those genes.


```
datExpr = list()

Expr = colnames(WGCNA_data[["datExpr"]])
Expr = Expr[which(Expr %in% my.ortho[,1])]

rownames(my.ortho) = my.ortho[,1]
Expr = my.ortho[Expr,]

for (t in 1:length(Wtest)) {
  Expr=Expr[which(Expr[,my.ortho.sp[t]] %in% rownames(Wtest[[t]]@assays$RNA@counts)),]
}

Expr=rownames(Expr)

for (t in 1:length(Wtest)) {
  x = my.ortho[,my.ortho.sp[t]][which(my.ortho[,1] %in% Expr)]
  datExpr[[t]] = t(as.matrix( Wtest[[t]]@assays$RNA@counts[x,] ))
  colnames(datExpr[[t]]) = my.ortho[,1][match(colnames(datExpr[[t]]), my.ortho[,my.ortho.sp[t]])]
}  

nonex=c()
for (i in datExpr) {
  nonex = c(nonex,which(!goodGenes(i)))
}
```


```
  ..Excluding 103 genes from the calculation due to too many missing samples or zero variance.
  ..Excluding 64 genes from the calculation due to too many missing samples or zero variance.
  ..Excluding 35 genes from the calculation due to too many missing samples or zero variance.
  ..Excluding 13 genes from the calculation due to too many missing samples or zero variance.
  ..Excluding 1 genes from the calculation due to too many missing samples or zero variance.
  ..Excluding 202 genes from the calculation due to too many missing samples or zero variance.
  ..Excluding 166 genes from the calculation due to too many missing samples or zero variance.
  ..Excluding 104 genes from the calculation due to too many missing samples or zero variance.
  ..Excluding 248 genes from the calculation due to too many missing samples or zero variance.
  ..Excluding 45 genes from the calculation due to too many missing samples or zero variance.
  ..Excluding 120 genes from the calculation due to too many missing samples or zero variance.
```


```
# nonex = c( nonex, Expr[which(!Expr %in% rownames(Wtest[[1]]@assays$RNA@counts))] )

nonex = unique(nonex)

Expr = colnames(datExpr[[1]])[-nonex]

for (t in 1:length(datExpr)) {
  datExpr[[t]] = datExpr[[t]][,Expr]
}
```


To make the actual test, we take the data from the WGCNA list, and the color labels from the dynamicMods vector. These will serve as reference to test the other networks against. Here we can see what’s the fraction of genes, from each module, that are survive the cur and are present in the comparison. We also take the expression data from the test sets, and make an object to pass to the testing function.


```
# Here we asign the reference data
refdat = WGCNA_data[["datExpr"]][,Expr]

# The reference colors
dynamicColors = labels2colors( WGCNA_data[["dynamicMods"]] )
names(dynamicColors) = colnames(WGCNA_data[["datExpr"]])
refcol = dynamicColors[Expr]

# Objects with both the reference and the test datasets
multiExpr = list(Reference = list(data = refdat))

for (t in 1:length(datExpr)) {
  multiExpr[[t+1]] = list(data = datExpr[[t]])
}

names(multiExpr) = c("Reference", Wtest_names)

multiColor = list(Reference = refcol)

my.percent = data.frame(percent = table(refcol)/table(labels2colors( WGCNA_data[["dynamicMods"]] )))

ggplot(my.percent, aes(x=percent.refcol, y=percent.Freq,fill =percent.refcol)) +
    geom_point( size =3.5, shape=21 ) + 
    scale_fill_manual(values = as.character(my.percent$percent.refcol)) + theme_minimal() +
    theme(legend.position ="none", text = element_text(size=15), axis.text.x = element_text(angle=45, hjust=1)) +
    labs(title = "Fraction of genes 1-to-1 orthologs,\nand present in test samples", x="Module", y="Fraction") +
    ylim(0, 1)
```


```
# The actual test

my.dummy=capture.output({
  mp = modulePreservation(multiExpr,
                          multiColor,
                          corFnc = "bicor",
                          maxGoldModuleSize = 300,
                          networkType = "signed",
                          referenceNetworks = 1,
                          nPermutations = 20,
                          randomSeed = 42,
                          quickCor = 0,
                          verbose = 0)
})
```


##### Results

We plot some results from the test, first the quality observed values against the preservation observed values, and then the quality Zscore against the preservation Zscore. These should MORE OR less correlate.


```
# To plot the results of the preservation tests

# Make lists to keep our datasets
statsObs = list()
statsZ = list()

# For each of the test samples, we make a data frame with the observed and Zsummary statisticals
for (test in 1:length(datExpr)) {
  statsObs[[test]] = cbind(mp$quality$observed[[1]][[test+1]],
                             mp$preservation$observed[[1]][[test+1]][,-1],
                             module = rownames(mp$quality$observed[[1]][[test+1]]))
  statsZ[[test]] = cbind(mp$quality$Z[[1]][[test+1]],
                           mp$preservation$Z[[1]][[test+1]][-1],
                           module = rownames(mp$quality$Z[[1]][[test+1]]))
}

# Compare preservation to quality:

# Make a list for the plotting
pq=list()
i=0

# Plot the preservations against the observed qualities. They should more or less correlate
for (test in 1:length(datExpr)) {
  plotData = data.frame(cbind(statsObs[[test]][, c("medianRank.pres", "medianRank.qual", "module")],
             signif(statsZ[[test]][, c("Zsummary.pres", "Zsummary.qual")], 2)))
  i=i+1
  pq[[i]] = ggplot(plotData, aes(x=medianRank.qual, y=medianRank.pres, label=module, fill=module)) + 
    geom_point(size=3.5, shape=21) +
    geom_text(nudge_x = 0.7, hjust=0, size=2.5, check_overlap = T) +
    scale_fill_manual(values= as.character(plotData$module)) + 
    theme_classic() + theme(legend.position="none") +
    scale_x_continuous(expand = c(0.1, 0)) +
    labs(title=paste0("Median rank in ", Wtest_names[test]), x ="Quality", y = "Preservation")
  i=i+1
  pq[[i]] = ggplot(plotData, aes(x=Zsummary.qual, y=Zsummary.pres, label=module, fill=module)) + 
    geom_point(size=3.5, shape=21) +
    geom_text(nudge_x = 0.7, hjust=0, size=2.5, check_overlap = T) +
    scale_fill_manual(values= as.character(plotData$module)) + 
    theme_classic() + theme(legend.position="none") +
    scale_x_continuous(expand = c(0.2, 0)) +
    labs(title=paste0("Zscore in ", Wtest_names[test]), x ="Quality", y = "Preservation")
}

grid.arrange(grobs=pq, ncol=2)
```


Now we can look at the actual preservation scores. We have two metrics, Zcore is useful in to compare against a threshold, and Median Rank which can be used to compare across modules:

- Zscores < 2 mean that there is no evidence of preservation of that module in that sample
- 2 < Zscores < 10 mean that there is moderate evidence of conservation of the module in that sample
- Zscores > 10 mean that the module is very well preserved in the sample
- Median Ranks are independent of module size, the smaller the Rank, the higher the preservation


```
# leave grey and gold modules out
modColors = rownames(mp$preservation$observed[[1]][[2]])
plotMods = !(modColors %in% c("grey", "gold"));

# The variable where we keep the data to plot
plotData = data.frame(Rank = numeric(), Zsum= numeric(), Size = numeric(), Cols = character(), Sample = character())
p=list()

# Fill in with data from the samples
for (test in 1:length(datExpr)) {
  plotData = rbind(plotData,data.frame(Rank = mp$preservation$observed[[1]][[test+1]][, 2],
                                       Zsum= mp$preservation$Z[[1]][[test+1]][, 2],
                                       Size = mp$preservation$observed[[1]][[test+1]][, 1],
                                       Cols = rownames(mp$preservation$observed[[1]][[test+1]]),
                                       Sample = rep(Wtest_names[test],nrow(mp$preservation$observed[[1]][[test+1]]))))
}

# We kick out the golden and grey modules
plotData = plotData[rep(plotMods, length(datExpr)),]


my.alpha = rep(1,dim(plotData)[1])
my.alpha[which(plotData$Zsum < 2)] = 0.75
plotData$alpha = my.alpha

# The plotting of the Zsummary and Median Rank, against the module size
p=list()

my.plotData = plotData
my.plotData$Sample = factor(my.plotData$Sample, levels = Wtest_names)
my.plotData$mylabels = my.plotData$Cols
my.plotData$mylabels[which(plotData$Zsum < 2)] = NA

x=0

if (class(my.grouping) == "list") {
  for (i in 1:length(my.grouping)) {
    plotData =my.plotData[which(my.plotData$Sample %in% Wtest_names[my.grouping[[i]]]),]

    if (length(my.grouping[[i]])>5) {
      my.gp = geom_point(aes(color=Cols, shape=Sample), size=3.5)
      my.scfm = scale_color_manual(values= as.character(unique(plotData$Cols)))
      my.ssm = scale_shape_manual(name= "Sample:",values= c(0:25), labels=Wtest_names[my.grouping[[i]]])
      my.gpa = geom_point(aes(color=Cols, shape=Sample, alpha=alpha), size=3.5)
    } else{
      my.gp = geom_point(aes(fill=Cols, shape=Sample), size=3.5)
      my.scfm = scale_fill_manual(values= as.character(unique(plotData$Cols)))
      my.ssm = scale_shape_manual(name= "Sample:",values= c(21:25), labels=Wtest_names[my.grouping[[i]]])
      my.gpa = geom_point(aes(fill=Cols, shape=Sample, alpha=alpha), size=3.5)
    }
  
  x=x+1
  
  p[[x]] = ggplot(plotData, aes(x=Size, y=Zsum, label=Cols)) + 
    geom_hline(yintercept = c(2,10), linetype="dashed", color=c("red", "limegreen")) +
    my.gp +
    geom_text(nudge_x = 0.08, hjust=0, size=2.5, check_overlap = T) +
    my.scfm + 
    my.ssm +
    theme_classic() + theme(legend.position="none") +
    scale_x_continuous(trans='log2', expand = c(0.1, 0)) +
    labs(x ="Module size", y = "Zsummary")
    
  x=x+1
  
  p[[x]] = ggplot(plotData, aes(x=Size, y=Rank, label=mylabels)) + 
    my.gpa +
    geom_text(nudge_x = 0.08, hjust=0, size=2.5, check_overlap = T) +
    my.scfm + 
    my.ssm +
    theme_classic() + theme(legend.text = element_text(size = 8), legend.title = element_text(size=10)) +
    guides(fill = FALSE, alpha = F, color=F) +
    scale_x_continuous(trans='log2', expand = c(0.1, 0)) +
    scale_y_continuous(trans = "reverse", expand = c(0.1,0)) +
    labs(x ="Module size", y = "Median Rank")
  
  }
} else {
  plotData = my.plotData
  if (length(levels(my.plotData))>5) {
      my.gp = geom_point(aes(color=Cols, shape=Sample), size=3.5)
      my.scfm = scale_color_manual(values= as.character(unique(plotData$Cols)))
      my.ssm = scale_shape_manual(name= "Sample:",values= c(0:25), labels=Wtest_names[my.grouping[[i]]])
      my.gpa = geom_point(aes(color=Cols, shape=Sample, alpha=alpha), size=3.5)
    } else{
      my.gp = geom_point(aes(fill=Cols, shape=Sample), size=3.5)
      my.scfm = scale_fill_manual(values= as.character(unique(plotData$Cols)))
      my.ssm = scale_shape_manual(name= "Sample:",values= c(21:25), labels=Wtest_names[my.grouping[[i]]])
      my.gpa = geom_point(aes(fill=Cols, shape=Sample, alpha=alpha), size=3.5)
    }
  p[[1]] = ggplot(plotData, aes(x=Size, y=Zsum, label=Cols)) + 
    geom_hline(yintercept = c(2,10), linetype="dashed", color=c("red", "limegreen")) +
    my.gp +
    geom_text(nudge_x = 0.08, hjust=0, size=2.5, check_overlap = T) +
    my.scfm + 
    my.ssm +
    theme_classic() + theme(legend.position="none") +
    scale_x_continuous(trans='log2', expand = c(0.1, 0)) +
    labs(x ="Module size", y = "Zsummary")
  
  p[[2]] = ggplot(plotData, aes(x=Size, y=Rank, label=mylabels)) + 
    my.gpa +
    geom_text(nudge_x = 0.08, hjust=0, size=2.5, check_overlap = T) +
    my.scfm + 
    my.ssm +
    theme_classic() + theme(legend.text = element_text(size = 8), legend.title = element_text(size=10)) +
    guides(fill = FALSE, alpha = F, color=F) +
    scale_x_continuous(trans='log2', expand = c(0.1, 0)) +
    scale_y_continuous(trans = "reverse", expand = c(0.1,0)) +
    labs(x ="Module size", y = "Median Rank")
}

grid.arrange(grobs=p, ncol=2, top="Preservation Zsummary and Median Rank")
```


But out of the different metrics from the module comparison, what is actually preserved? For this we can look at the density and connectivity preservations. **Density** tests if the module nodes remain highly connected in the test networks **Connectivity** tests if the connectivity pattern between the nodes is similar in the test networks


```
# leave grey and gold modules out
plotMods = !(modColors %in% c("grey", "gold"));

# The variable where we keep the data to plot
plotData = data.frame(Density = numeric(), Connectivity = numeric(), Size = numeric(), Cols = character(), Sample = character())
p=list()

# Fill in with data from the samples
for (test in 1:length(datExpr)) {
  plotData = rbind(plotData,data.frame(Density = mp$preservation$Z[[1]][[test+1]][, 3],
                                       Connectivity = mp$preservation$Z[[1]][[test+1]][, 4],
                                       Size = mp$preservation$observed[[1]][[test+1]][, 1],
                                       Cols = rownames(mp$preservation$observed[[1]][[test+1]]),
                                       Sample = rep(Wtest_names[test],nrow(mp$preservation$observed[[1]][[test+1]]))))
}

# We kick out the golden and grey modules
plotData = plotData[rep(plotMods, length(datExpr)),]

# The plotting of the Zsummary and Median Rank, against the module size
p=list()

my.lim = (max(plotData[,1:2])-min(plotData[,1:2]))*0.025
my.lim = c(min(plotData[,1:2])-my.lim, max(plotData[,1:2])+my.lim)

p=list()

my.plotData = plotData
my.plotData$Sample = factor(my.plotData$Sample, levels = Wtest_names)
my.plotData$mylabels = my.plotData$Cols
my.plotData$mylabels[which(plotData$Zsum < 2)] = NA

x=0

if (class(my.grouping) == "list") {
  for (i in 1:length(my.grouping)) {
    plotData =my.plotData[which(my.plotData$Sample %in% Wtest_names[my.grouping[[i]]]),]

    if (length(my.grouping[[i]])>5) {
      my.gp = geom_point(aes(color=Cols, shape=Sample), size=3.5)
      my.scfm = scale_color_manual(values= as.character(unique(plotData$Cols)))
      my.ssm = scale_shape_manual(name= "Sample:",values= c(0:25), labels=Wtest_names[my.grouping[[i]]])
    } else{
      my.gp = geom_point(aes(fill=Cols, shape=Sample), size=3.5)
      my.scfm = scale_fill_manual(values= as.character(unique(plotData$Cols)))
      my.ssm = scale_shape_manual(name= "Sample:",values= c(21:25), labels=Wtest_names[my.grouping[[i]]])
    }
  
  x=x+1
  
  p[[x]] = ggplot(plotData, aes(x=Size, y=Density, label=Cols)) + 
    geom_hline(yintercept = c(2,10), linetype="dashed", color=c("red", "limegreen")) +
    my.gp +
    geom_text(nudge_x = 0.08, hjust=0, size=2.5, check_overlap = T) +
    my.scfm + 
    my.ssm +
    theme_classic() + theme(legend.position="none") +
    scale_x_continuous(trans='log2', expand = c(0.1, 0)) +
    labs(x ="Module size", y = "Density") + ylim(my.lim)
    
  x=x+1
  
  p[[x]] = p[[x]] = ggplot(plotData, aes(x=Size, y=Connectivity, label=Cols)) + 
    geom_hline(yintercept = c(2,10), linetype="dashed", color=c("red", "limegreen")) +
    my.gp +
    geom_text(nudge_x = 0.08, hjust=0, size=2.5, check_overlap = T) +
    my.scfm + 
    my.ssm +
    theme_classic() +
    scale_x_continuous(trans='log2', expand = c(0.1, 0)) +
    labs(x ="Module size", y = "Connectivity") + theme(legend.text = element_text(size = 8), legend.title = element_text(size=10)) +
    guides(fill = FALSE, color=F) + ylim(my.lim)
  
  }
} else {
  plotData = my.plotData
  if (length(levels(my.plotData))>5) {
      my.gp = geom_point(aes(color=Cols, shape=Sample), size=3.5)
      my.scfm = scale_color_manual(values= as.character(unique(plotData$Cols)))
      my.ssm = scale_shape_manual(name= "Sample:",values= c(0:25), labels=Wtest_names[my.grouping[[i]]])
    } else{
      my.gp = geom_point(aes(fill=Cols, shape=Sample), size=3.5)
      my.scfm = scale_fill_manual(values= as.character(unique(plotData$Cols)))
      my.ssm = scale_shape_manual(name= "Sample:",values= c(21:25), labels=Wtest_names[my.grouping[[i]]])
    }
  p[[1]] = ggplot(plotData, aes(x=Size, y=Density, label=Cols)) + 
    geom_hline(yintercept = c(2,10), linetype="dashed", color=c("red", "limegreen")) +
    my.gp +
    geom_text(nudge_x = 0.08, hjust=0, size=2.5, check_overlap = T) +
    my.scfm + 
    my.ssm +
    theme_classic() + theme(legend.position="none") +
    scale_x_continuous(trans='log2', expand = c(0.1, 0)) +
    labs(x ="Module size", y = "Density") + ylim(my.lim)
  
  p[[2]] = ggplot(plotData, aes(x=Size, y=Connectivity, label=Cols)) + 
    geom_hline(yintercept = c(2,10), linetype="dashed", color=c("red", "limegreen")) +
    my.gp +
    geom_text(nudge_x = 0.08, hjust=0, size=2.5, check_overlap = T) +
    my.scfm + 
    my.ssm +
    theme_classic() +
    scale_x_continuous(trans='log2', expand = c(0.1, 0)) +
    labs(x ="Module size", y = "Connectivity") + theme(legend.text = element_text(size = 8), legend.title = element_text(size=10)) +
    guides(fill = FALSE, color=F) + ylim(my.lim)
  
}

grid.arrange(grobs=p, ncol=2, top="Preservation of Density and Connectivity")
```


Saving the preservation data under ./data/E15\_modulepreservation\_231220.rds


```
saveRDS(mp, paste0("./data/",project_name,"_modulepreservation_",format(Sys.Date(), "%d%m%y"),".rds"))
```
